# TRIP13 localizes to synapsed chromosomes and functions as a dosage-sensitive regulator of meiosis

**DOI:** 10.1101/2023.09.25.559355

**Authors:** Jessica Y. Chotiner, N. Adrian Leu, Fang Yang, Isabella G. Cossu, Yongjuan Guan, Huijuan Lin, P. Jeremy Wang

## Abstract

Meiotic progression requires coordinated assembly and disassembly of protein complexes involved in chromosome synapsis and meiotic recombination. The AAA+ ATPase TRIP13 and its orthologue Pch2 are instrumental in remodeling HORMA domain proteins. Meiosis-specific HORMAD proteins are associated with unsynapsed chromosome axes but depleted from the synaptonemal complex (SC) of synapsed chromosome homologues. Here we report that TRIP13 localizes to the synapsed SC in early pachytene spermatocytes and to telomeres throughout meiotic prophase I. Loss of TRIP13 leads to meiotic arrest and thus sterility in both sexes. *Trip13*-null meiocytes exhibit abnormal persistence of HORMAD1 and HOMRAD2 on synapsed SC and chromosome asynapsis that preferentially affects XY and centromeric ends. These findings confirm the previously reported phenotypes of the *Trip13* hypomorph alleles. *Trip13* heterozygous (*Trip13*^+/-^) mice also exhibit meiotic defects that are less severe than the *Trip13*-null mice, showing that TRIP13 is a dosage-sensitive regulator of meiosis. Localization of TRIP13 to the synapsed SC is independent of SC axial element proteins such as REC8 and SYCP2/SYCP3. The N- or C-terminal FLAG-tagged TRIP13 proteins are functional and recapitulate the localization of native TRIP13 to SC and telomeres in knockin mice. Therefore, the evolutionarily conserved localization of TRIP13/Pch2 to the synapsed chromosomes provides an explanation for dissociation of HORMA domain proteins upon chromosome synapsis in diverse organisms.

## Introduction

Meiosis is a specialized cell division program that generates haploid gametes from diploid germ cells. During meiotic prophase I, homologous chromosomes undergo paring, synapsis, and recombination (Handel and Schimenti, 2010; Zickler and Kleckner, 2015). Chromosome synapsis requires the formation of the synaptonemal complex (SC). The SC is a proteinaceous structure comprised of two lateral/axial elements, transverse filaments, and a central element (Page and Hawley, 2004). Meiotic recombination begins with generation of DNA double strand breaks (DSBs) and ends with crossover formation through repair of DSBs (Hunter, 2015). Chromosome synapsis and meiotic recombination are interdependent in many species including yeast and mouse. Meiotic recombination not only increases genetic diversity in gametes at the organism level but also is essential for faithful chromosome segregation during the first meiotic cell division. Therefore, defects in meiosis are leading causes of aneuploidy, birth defect, infertility, and pregnancy loss.

The progression and completion of synapsis and recombination are monitored by a surveillance mechanism called meiotic checkpoint (Roeder and Bailis, 2000). Meiotic checkpoint proteins were extensively studied in yeast. Many meiotic processes are highly conserved, and studies of meiosis in yeast have been foundational for our understanding of mammalian meiosis. In yeast, Pch2, an AAA+ ATPase, is a checkpoint protein, because it is necessary for pachytene arrest in the absence of the SC protein Zip1 or the meiosis-specific DSB repair protein Dmc1 (Herruzo et al., 2021; San-Segundo and Roeder, 1999). Pch2 is a hexameric ring ATPase and remodels the HORMA (<u>Ho</u>p1, <u>R</u>ev7, and <u>MA</u>D2) domain protein Hop1 in a nucleotide-dependent manner (Chen et al., 2014). While Pch2 is mainly located in the nucleolus, Pch2 also localizes to the synapsed SC as foci at the pachytene stage and is essential for removing Hop1 from chromosome axes (Chen et al., 2014; Joshi et al., 2009; San-Segundo and Roeder, 1999). Mechanistically, Pch2 promotes phosphorylation of Hop1, which activates the Mek1 kinase and the subsequent checkpoint cascade (Herruzo et al., 2016; Raina and Vader, 2020).

TRIP13, the mammalian orthologue of yeast Pch2, is essential for completion of meiotic recombination in mouse (Li and Schimenti, 2007; Roig et al., 2010). Mutations in human *TRIP13* predispose to Wilms tumor in children or cause infertility in women (Yost et al., 2017; Zhang et al., 2020). HORMAD1 and HORMAD2, mammalian meiosis-specific HORMA domain proteins, localize to chromosome axis at leptotene and zygotene stages but are depleted from synapsed chromosomes except the largely unsynapsed XY chromosomes at the pachytene stage (Fukuda et al., 2010; Xu et al., 2019). TRIP13 facilitates removal of HORMAD proteins from synapsed chromosome axis (Roig et al., 2010; Wojtasz et al., 2009). SKP1, an essential component of the SCF (Skp1-Cullin 1-F-box protein) ubiquitin E3 ligase, is required not only for depleting HORMAD1 and HORMAD2 from synapsed chromosome axes but also for restricting the accumulation of HORMAD proteins on unsynapsed axes (Guan et al., 2020; Guan et al., 2022). Therefore, reorganization of HORMA domain proteins on meiotic chromosome axes is regulated by both TRIP13 and SKP1. As a conserved mechanism, Pch2/TRIP13 remodels HORMA domain proteins through engagement of their N-terminal regions (Prince and Martinez-Perez, 2022). MAD2, a spindle assembly checkpoint HORMA domain protein, exists in two fold states: closed and open (Luo et al., 2004; Sironi et al., 2002). In the closed state, the so-called safety belt of the HORMA domain wraps around the HORMAD protein-binding motif “closure motif” in the partner protein and thus traps the partner. In the open state, TRIP13 engagement changes conformation of the HORMA domain, resulting in foldback of the safety belt on the closure motif-binding region and thus release of its partner protein. The safety belt mechanism turns out to be a shared feature of HORMA domain proteins (Kim et al., 2014; West et al., 2019; West et al., 2018; Ye et al., 2017).

Unlike yeast Pch2, TRIP13 does not appear to function in the meiotic checkpoint in mouse (Pacheco et al., 2015). Interestingly, HORMAD2 functions as a meiotic checkpoint protein for surveillance of meiotic defects in female meiosis (Kogo et al., 2012a; Rinaldi et al., 2017; Wojtasz et al., 2012). HORMAD2-deficient males are sterile but females are fertile. HORMAD2 deficiency rescues the oocyte loss in *Spo11*-null or *Trip13* mutant females but not in *Dmc1*-null females, suggesting that HORMAD2 is a component of the meiotic checkpoint in females. HORMAD1 is required for fertility in both sexes (Kogo et al., 2012b; Shin et al., 2010; Stanzione et al., 2016). Both HORMAD1 and HORMAD2 localize prominently to the XY chromosome axes in pachytene spermatocytes and regulate meiotic sex chromatin inactivation (MSCI) (Kogo et al., 2012a; Shin et al., 2010; Wojtasz et al., 2012).

Studies of mice with hypomorphic *Trip13* mutations show that TRIP13 is required for meiotic progression (Li and Schimenti, 2007; Roig et al., 2010). Two different hypomorphic *Trip13* mouse lines have been studied: one “moderate” allele that exhibited defective meiotic recombination and one “severe” allele that displayed defects in meiotic recombination and chromosome synapsis. The variance in phenotype between the hypomorphic mouse lines indicates that even partial depletion of *Trip13* can interrupt meiosis. We sought to further investigate the meiotic function of TRIP13. By immunofluorescence, we found that TRIP13 localized to the synaptonemal complex in early pachytene spermatocytes and to telomeres throughout meiotic prophase I. The TRIP13 localization pattern on meiotic chromosomes was further confirmed by analysis of FLAG-tagged TRIP13 in two knockin mouse lines. To determine the effect of complete loss of TRIP13, we characterized *Trip13*-null mice. While the *Trip13*-null phenotype was largely comparable to the phenotype of the severe *Trip13* hypomorphic allele, we found that TRIP13 is a dosage-sensitive regulator of meiosis. Finally, our results show that the localization of TRIP13 to the SC is independent of axial element proteins.

## Results

### TRIP13 localizes to meiotic chromosomes in prophase I spermatocytes

We assessed the expression of TRIP13 in a panel of adult mouse tissues. TRIP13 was primarily expressed in testis but detected at low levels in ovary and liver (Figure 1A). In the testis, TRIP13 was prominent in the cytoplasm of primary spermatocytes, especially leptotene and zygotene cells (Figure 1B). Given the expression of TRIP13 in spermatocytes, we examined its localization on meiotic chromosomes by immunostaining of surface spread nuclei of prophase I spermatocytes (Figure 1C). Previously, TRIP13 was reported to localize to telomeres in spermatocytes (Gomez et al., 2019). Indeed, we found that TRIP13 localized to telomeres from leptotene to diplotene cells (Figure 1C). Strikingly, we found that, in addition to telomeres, TRIP13 localized to the synaptonemal complex (SC) in early pachytene spermatocytes but disappeared from the SC in the mid to late pachytene spermatocytes (Figure 1C). The synaptonemal complex consists of two lateral elements and one central element. The central region is comprised of the transverse filament protein SYCP1 and central element proteins, which appear upon synapsis. Given the novel localization of TRIP13 to the SC, we performed confocal immunofluorescent microscopy with super-resolution deconvolution to determine the location of TRIP13 within the SC. In contrast with the lateral element marker SYCP3, TRIP13 localized to the SC central region (Figure 1D). We examined the localization of TRIP13 in oocytes. TRIP13 localized strongly to meiotic telomeres in all stages of prophase I, but not as filaments on the SC (Figure 1 - figure supplement 1). These findings are consistent with the genetic requirement of TRIP13 in meiosis in both sexes (Li and Schimenti, 2007; Roig et al., 2010).

**Figure 1.**
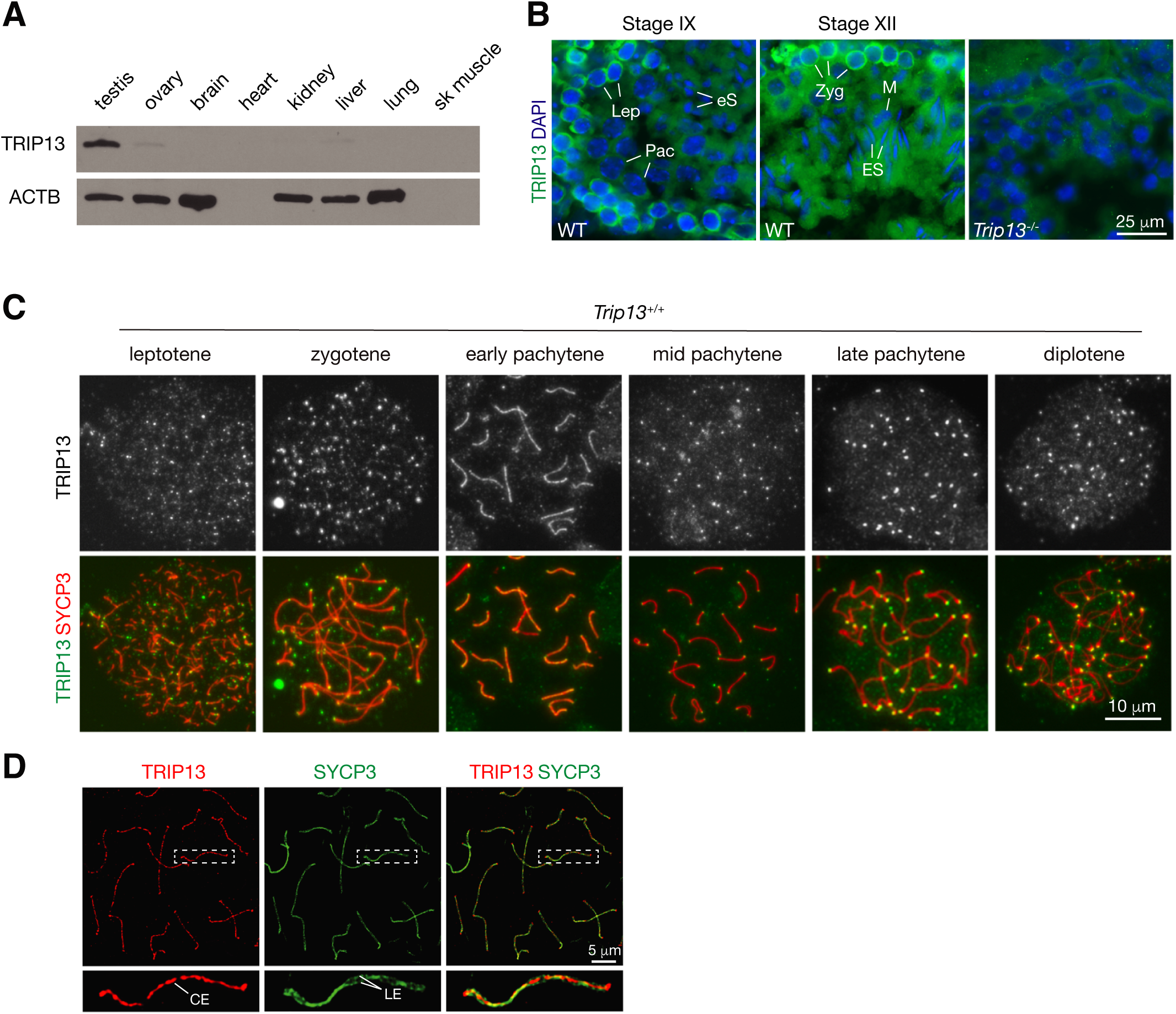
TRIP13 localizes to the synaptonemal complex and telomeres in spermatocytes. (A) Western blot analysis of TRIP13 in adult mouse tissues. Heart and skeletal muscle lack ACTB. (B) Immunofluorescence of TRIP13 in sections of 3-month-old wild type and *Trip13*^-/-^ testes. Lep, leptotene; Zyg, zygotene; Pac, pachytene; eS, elongating spermatids; ES, elongated spermatids. (C) Immunofluorescence of TRIP13 in spread nuclei of spermatocytes from wild type P20 testes. (D) Super-resolution localization of TRIP13 to the central element (CE) but not lateral element (LE) of the synaptonemal complex at early pachytene stage. The enlarged view of the boxed chromosome is shown at the bottom.

### Global loss of *Trip13* causes meiotic arrest

Two previous studies utilizing two *Trip13* hypomorphic (gene trap) alleles, one moderate and one severe, revealed a role for TRIP13 in meiotic recombination (Li and Schimenti, 2007; Roig et al., 2010). However, while the *Trip13* moderate allele showed mostly intact chromosomal synapsis, the severe allele displayed chromosomal unsynapsis preferentially at chromosome ends (Li and Schimenti, 2007; Roig et al., 2010). Upon close examination, we also found unsynapsed ends in spermatocytes with the moderate *Trip13* mutant (Guan et al., 2020). To rigorously ascertain the role of TRIP13 in meiosis, we generated *Trip13*-null mutants using frozen sperm from *Trip13*^+/-^ males with a knockout allele from the International Mouse Phenotyping Consortium (IMPC). The mouse *Trip13* gene consists of 13 exons. This new *Trip13* mutant allele harbors a deletion of a 24-kb region including all 13 exons and thus is expected to be null. We verified this deletion by PCR and sequencing. Interbreeding of *Trip13*^+/-^ mice produced fewer *Trip13*^-/-^ offspring than expected: *Trip13*^+/+^, 80; *Trip13*^+/-^, 220; *Trip13*^-/-^, 51 (χ2=27, *p*=0.0001). The body weight of 2-3 month-old males was not significantly different between wild type (24.3±2.8 g, n=5) and *Trip13*^-/-^ mice (22.8±1.7 g, n=5, *p*=0.3, Student’s *t*-Test). The *Trip13*^-/-^ testis was much smaller than *Trip13*^+/-^ testis (Figure 2A). Western blot analysis showed that TRIP13 was present in reduced abundance in *Trip13*^+/-^ testis and not detected in *Trip13*^-/-^ testis, demonstrating that this mutant allele is null (Figure 2B). The abundance of SYCP3, a component of the synaptonemal complex, was also reduced in *Trip13*^-/-^ testis, suggesting a partial depletion of meiotic cells (Figure 2B). The testis weight of adult *Trip13*^-/-^ males was reduced by 74% in comparison with the wild type males (Figure 2C). Adult *Trip13*^-/-^ males lacked sperm in the epididymis (Figure 2D). Histological analysis showed that while spermatocytes in all stages of meiosis were present in adult *Trip13*^+/+^ and *Trip13*^+/-^ testes, *Trip13*^-/-^ testis displayed complete meiotic arrest, evidenced by the presence of early spermatocytes and a lack of secondary spermatocytes, round spermatids, and mature sperm (Figure 2E). *Trip13*^-/-^ females were also sterile. The adult *Trip13*^-/-^ ovary was very small and showed a complete loss of oocytes (Figure 2 - figure supplement 1). These observations are similar to the meiotic arrest phenotype observed in the hypomorphic *Trip13* mouse mutants (Li and Schimenti, 2007; Roig et al., 2010).

**Figure 2.**
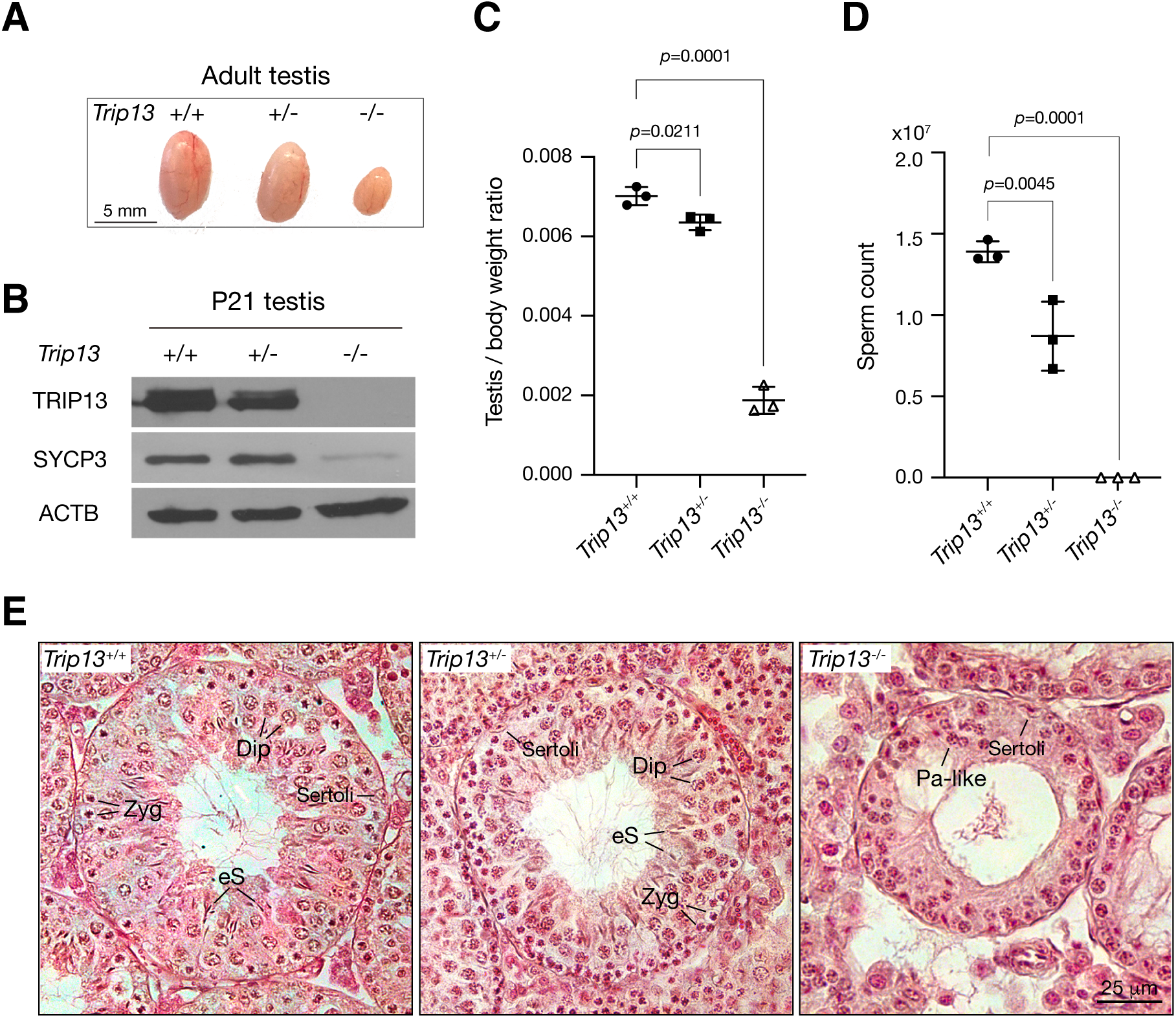
Loss of *Trip13* leads to meiotic arrest in males. (A) Image of testes from 2 to 3 month-old mice. (B) Western blot analysis of TRIP13 in P21 testes. SYCP3 serves as a meiosis-specific marker. ACTB serves as a loading control. (C) Testis to body weight ratio of 2 to 3 month-old mice. n = 3 males. Statistics, one-Way ANOVA. (D) Sperm count of 2 to 3 month-old *Trip13*^+/+^ and *Trip13*^+/-^ males. n = 3 males. Statistics, one-Way ANOVA. (E) Histological analysis of 2-month-old testes. Sertoli, Sertoli cell; Zyg, zygotene; Pa-like, pachytene-like; Dip, diplotene; eS, elongating spermatids.

### Defects in chromosomal synapsis in *Trip13*-deficient spermatocytes

We investigated the effect of *Trip13* deficiency on meiotic progression. The transverse filament/central element protein SYCP1 and the lateral element protein SYCP3 were used to determine the stage of prophase I spermatocytes. While the P20 wild type testis contained spermatocytes from leptotene through diplotene stages, about half of the *Trip13*^-/-^ spermatocytes were in the early pachytene stage and no cells were found at later stages of prophase I, showing meiotic blockade at the early pachytene stage in *Trip13*^-/-^ testes (Figure 3A). The early pachytene-like spermatocytes from *Trip13*^-/-^ testes contained unsynapsed chromosomal ends (Figure 3B). Mouse centromeres are telocentric. Co-staining with the centromere marker CREST showed that 94% of unsynapsed ends were centromeric ends (Figure 4A). While XY chromosomes were synapsed at the pseudoautosomal regions and thus were connected in wild type and *Trip13*^+/-^ pachytene spermatocytes, they were separate in *Trip13*^-/-^ pachytene-like spermatocytes (Figure 3B and Figure 3C). We found that SYCE1, a component of the SC central element, localized to the synapsed SC but not to unsynapsed regions in *Trip13*^-/-^ pachytene-like cells (Figure 3C). Confocal microscopy with super-resolution deconvolution confirmed that many homologous chromosomes in *Trip13*^-/-^ spermatocytes had split ends and some had regions of interstitial asynapsis (Figure 3D). We quantified these meiotic defects in spermatocytes: chromosomal asynapsis (Figure 3E), the number of asynapsed ends (Figure 3F), and the extent of XY asynapsis (Figure 3G). Previous studies also reported synaptic defects in spermatocytes from *Trip13* hypomorph mutants (Li and Schimenti, 2007; Roig et al., 2010). Here we found that global loss of *Trip13* caused similar defects in autosomal synapsis but a more severe defect in sex chromosome synapsis.

**Figure 3.**
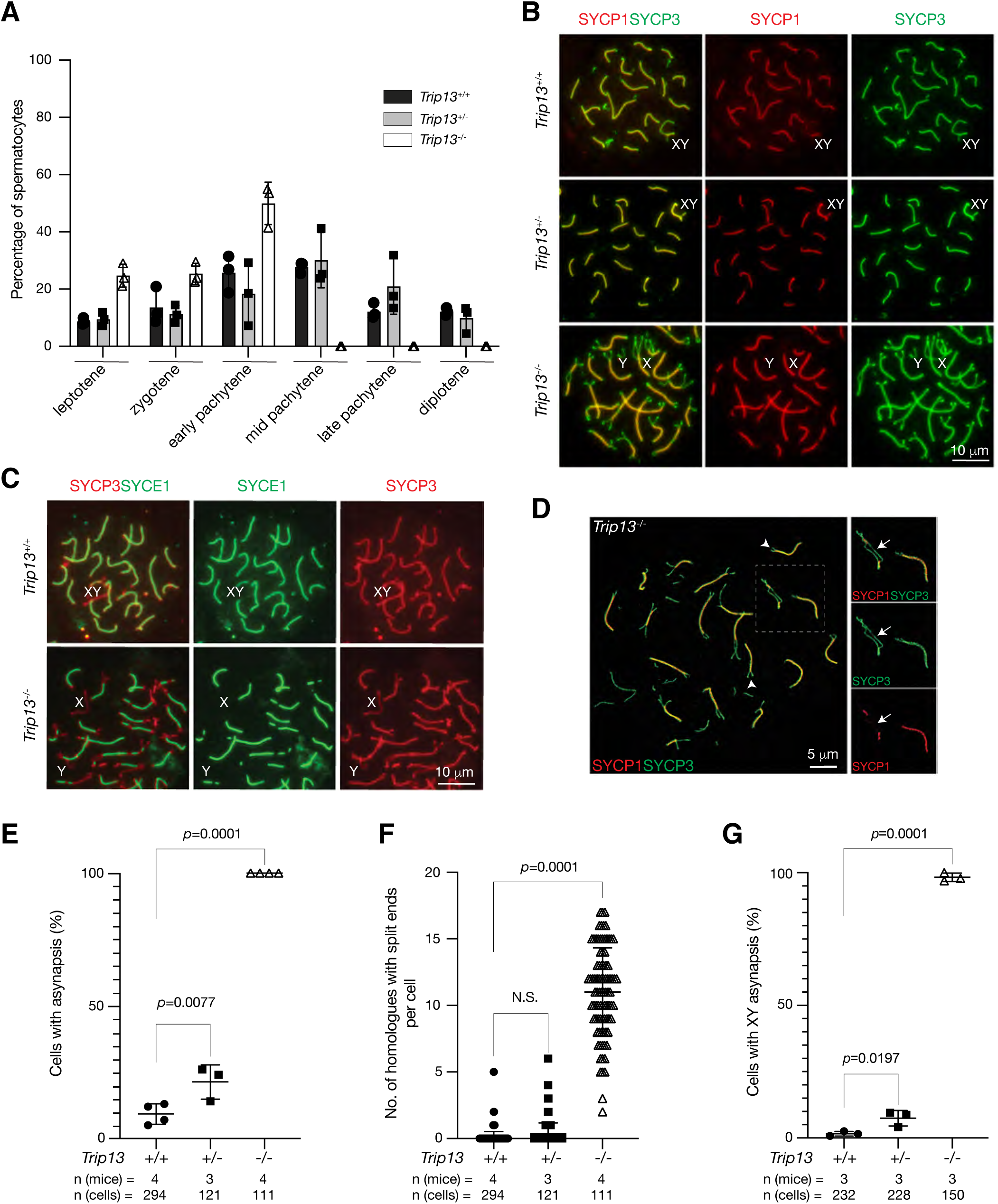
*Trip13* is required for chromosomal synapsis in males. (A) Composition of prophase I spermatocytes in P20 testes. Three males per genotype were analyzed by nuclear spread analysis. Total number of spermatocytes counted: *Trip13*^+/+^, 1038 cells; *Trip13*^+/-^, 803 cells; *Trip13*^-/-^, 406 cells. (B) Immunofluorescence of SYCP1 and SYCP3 in spread nuclei of pachytene spermatocytes from P20 testes. (C) Immunofluorescence of SYCE1 and SYCP3 in spread nuclei of pachytene spermatocytes from P20 testes. (D) Super-resolution confocal microscopy of a *Trip13*^-/-^ spermatocyte from P20 testis. Immunostaining was performed for SYCP1 and SYCP3. Arrowheads indicate end asynapsis. Arrow indicates interstitial asynapsis. (E) Percentage of early pachytene cells from P19-20 testes with asynapsed chromosomes across three genotypes. (F) Number of homologous chromosomes with end asynapsis per cell in P19-20 testes. (G) Percentage of early pachytene cells from P20 testes with asynapsed XY chromosomes per mouse. The *p* values are indicated in graphs. Statistics (E-G), one-way ANOVA.

**Figure 4.**
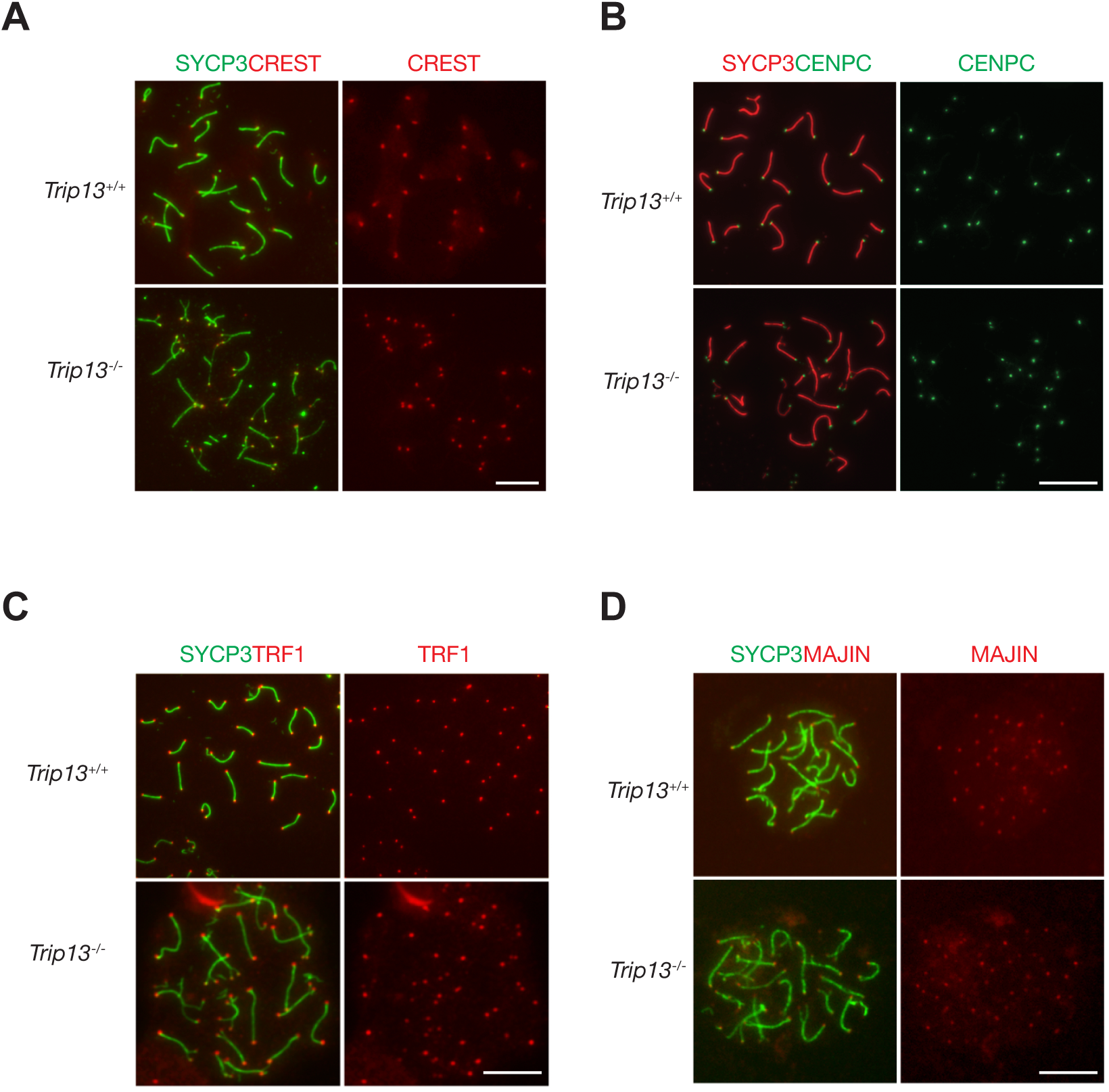
Normal localization of centromere and telomere markers in *Trip13*-deficient spermatocytes from juvenile mice. (A-D) Immunofluorescent analysis of centromere and telomere markers in *Trip13*^+/+^ and *Trip13*^-/-^ pachytene spermatocytes: CREST (A), CENPC (B), TRF1 (C), and MAJIN (D). SYCP3 labels the lateral elements of the synaptonemal complex. Scale bars, 10 µm.

### TRIP13 is a dosage-sensitive regulator of meiosis

Although *Trip13*^+/-^ males were fertile, their testis weight and sperm count were significantly reduced in comparison with wild type (Figure 2C-D). The percentage of spermatocytes with defects in synapsis was higher in *Trip13*^+/-^ males than wild type (Figure 3E). In addition, the percentage of spermatocytes with asynapsed XY chromosomes was significantly higher in *Trip13*^+/-^ males than wild type (Figure 3G). As defects in synapsis can activate the meiotic checkpoint and thus apoptosis of affected spermatocytes, the increased synaptic defects likely caused the decrease in sperm output in *Trip13*^+/-^ males. These results demonstrate that TRIP13 regulates meiosis in a dosage-dependent manner.

### Telomere and centromere proteins localize normally in *Trip13*-deficient spermatocytes

TRIP13 localizes to telomeres and loss of TRIP13 causes peri-centromeric/telomeric asynapsis. Therefore, we asked whether telomere or centromere proteins were affected in *Trip13*^-/-^ spermatocytes. Chromosome spreads were immunostained for two centromere markers, CREST and CENPC (Figure 4A-B). CENPC is essential for recruiting kinetochore proteins to the centromere (Klare et al., 2015; Kwon et al., 2007). CREST and CENPC still localized to centromeres in *Trip13*^-/-^ spermatocytes. Because many *Trip13*^-/-^ spermatocytes had split ends, two CREST or two CENPC foci were observed at the split ends. To assess the telomere, spermatocytes were immunostained for TRF1 and MAJIN (Figure 4C-D). In meiotic cells, telomeres contain canonical telomere proteins such as TRF1/TRF2 and a meiosis-specific complex (MAJIN, TERB1, TERB2, and SUN1) (Shibuya et al., 2015). TRF1 is a telomere protein expressed in both somatic and germ cells (Karlseder et al., 2003; Long et al., 2017; Shibuya et al., 2014). MAJIN is a component of the meiosis-specific telomere complex that is important for telomere attachment to the inner nuclear membrane (Shibuya et al., 2015). Both TRF1 and MAJIN localized to telomeres in *Trip13*^-/-^ spermatocytes (Figure 4C-D). As expected, two TRF1 or two MAJIN foci were observed at split ends of chromosomes in *Trip13*^-/-^ spermatocytes (Figure 4C-D). ANKRD31 and REC114 interact with each other and both localize to the pseudoautosomal region (PAR) of XY chromosomes in early pachytene spermatocytes to ensure XY recombination (Boekhout et al., 2019; Papanikos et al., 2019). In addition to two foci on autosomes, REC114 localized as one focus on the PAR in wild type pachytene spermatocytes. REC114 still formed foci (one per chromosome) on the unsynapsed X and Y chromosomes in *Trip13*^-/-^ pachytene-like spermatocytes (Figure 4 - figure supplement 1). Taken together, these results suggest that TRIP13 is not required for recruitment of these centromere or telomere proteins.

### TRIP13 is required to evict HORMAD1 and HORMAD2 from synapsed autosomes

In wild type meiotic cells, HORMAD1 and HORMAD2 localize to unsynapsed and desynapsed chromosomes (Fukuda et al., 2010; Kogo et al., 2012a; Shin et al., 2010; Wojtasz et al., 2012). Thus, HORMAD1 and HORMAD2 localized to the largely unsynapsed XY but not to synapsed autosomal SCs in wild type cells (Figure 5A-B). TRIP13 is essential for removing meiotic HORMADs from the chromosome axes, a function that is conserved in yeast, worms, and mammals (Wojtasz et al., 2009). We confirmed that HORMAD1 and HORMAD2 remained on the synapsed autosomes in *Trip13*^-/-^ pachytene cells (Figure 5A-B). HORMAD1 and HORMAD2 localize to the interior of the lateral elements in the SC (Xu et al., 2019). Confocal microscopy with super-resolution deconvolution revealed that HORMAD1 and HORMAD2 localized to the lateral elements of the synapsed autosomes in *Trip13*^-/-^ spermatocytes (Figure 5C-D). These results were consistent with accumulation of HORMAD1/2 in *Trip13* hypomorphic mutant spermatocytes (Roig et al., 2010; Wojtasz et al., 2009). Our analysis of *Trip13*-null mutant confirmed that TRIP13 is essential for HORMAD1/2 removal from the synapsed SCs.

**Figure 5.**
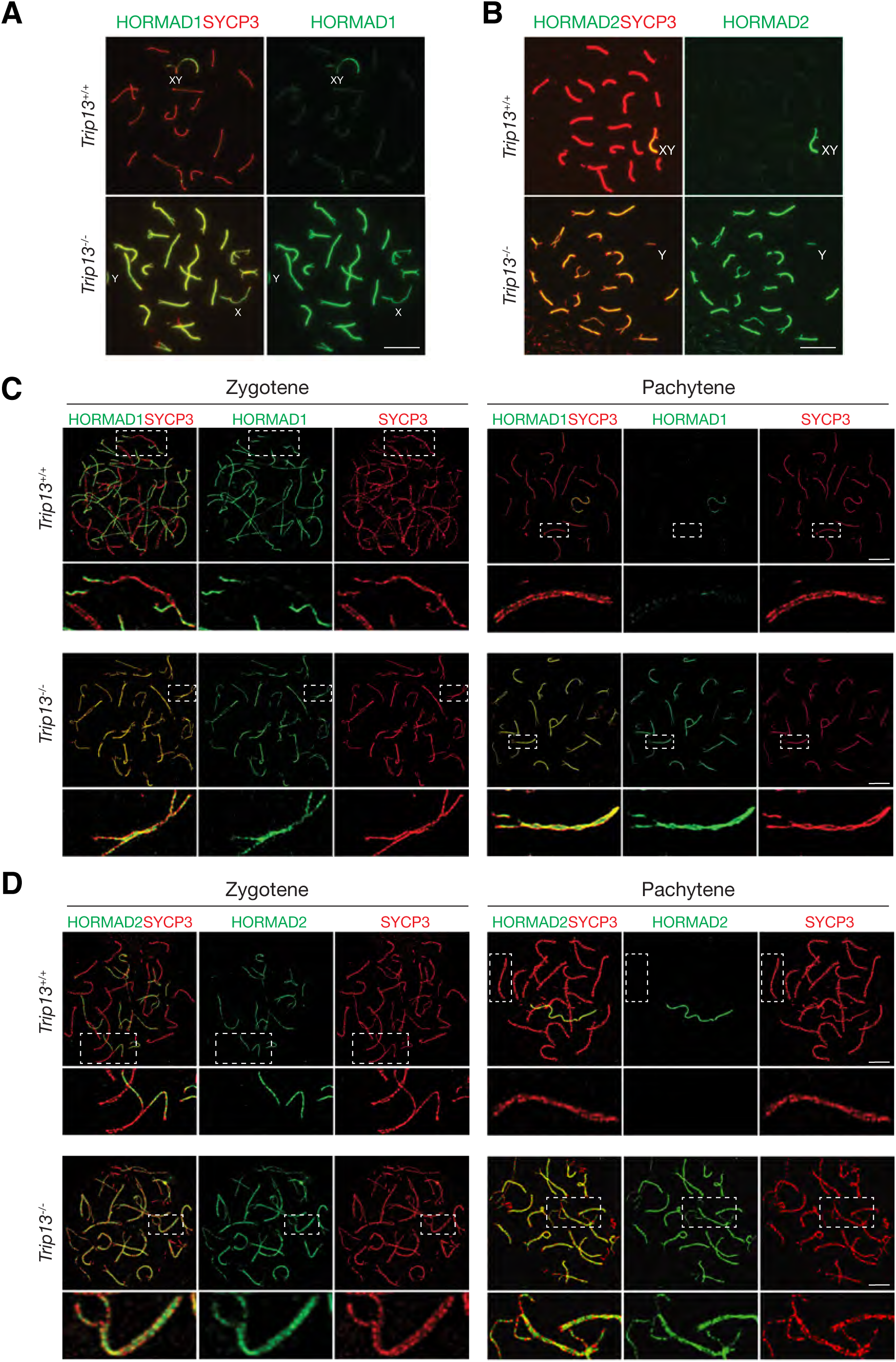
HORMAD1 and HORMAD2 accumulate on the lateral elements of synapsed autosomes in *Trip13*^-/-^ spermatocytes from juvenile (P19-21) mice. (A, B) Immunofluorescence of HORMAD1 (A) and HORMAD2 (B) in pachytene spermatocytes. (C, D) Super-resolution imaging of HORMAD1 and HORMAD2 in zygotene and pachytene spermatocytes. Enlarged views of the boxed chromosomes are shown below. Scale bars: 10 µm (A, B), 5 µm (C, D).

### Localization of TRIP13 to SC is independent of axial element components

TRIP13 localizes to the SC in early pachytene spermatocytes. We sought to address what proteins might recruit TRIP13 to the SC. We examined the role of HORMAD1, REC8, SYCP2, and SKP1 in TRIP13 localization in spermatocytes using respective knockout mice (Figure 6). We first examined the localization of TRIP13 in *Hormad1*^-/-^ spermatocytes. *Hormad1*^-/-^ spermatocytes exhibit limited synapsis (Daniel et al., 2011; Kogo et al., 2012b; Shin et al., 2010). In both wild type and *Hormad1*^-/-^ cells, TRIP13 localized to the synapsed regions and to the telomeres (Figure 6A). This result shows that HORMAD1 is not required for localization of TRIP13 to the SC.

**Figure 6.**
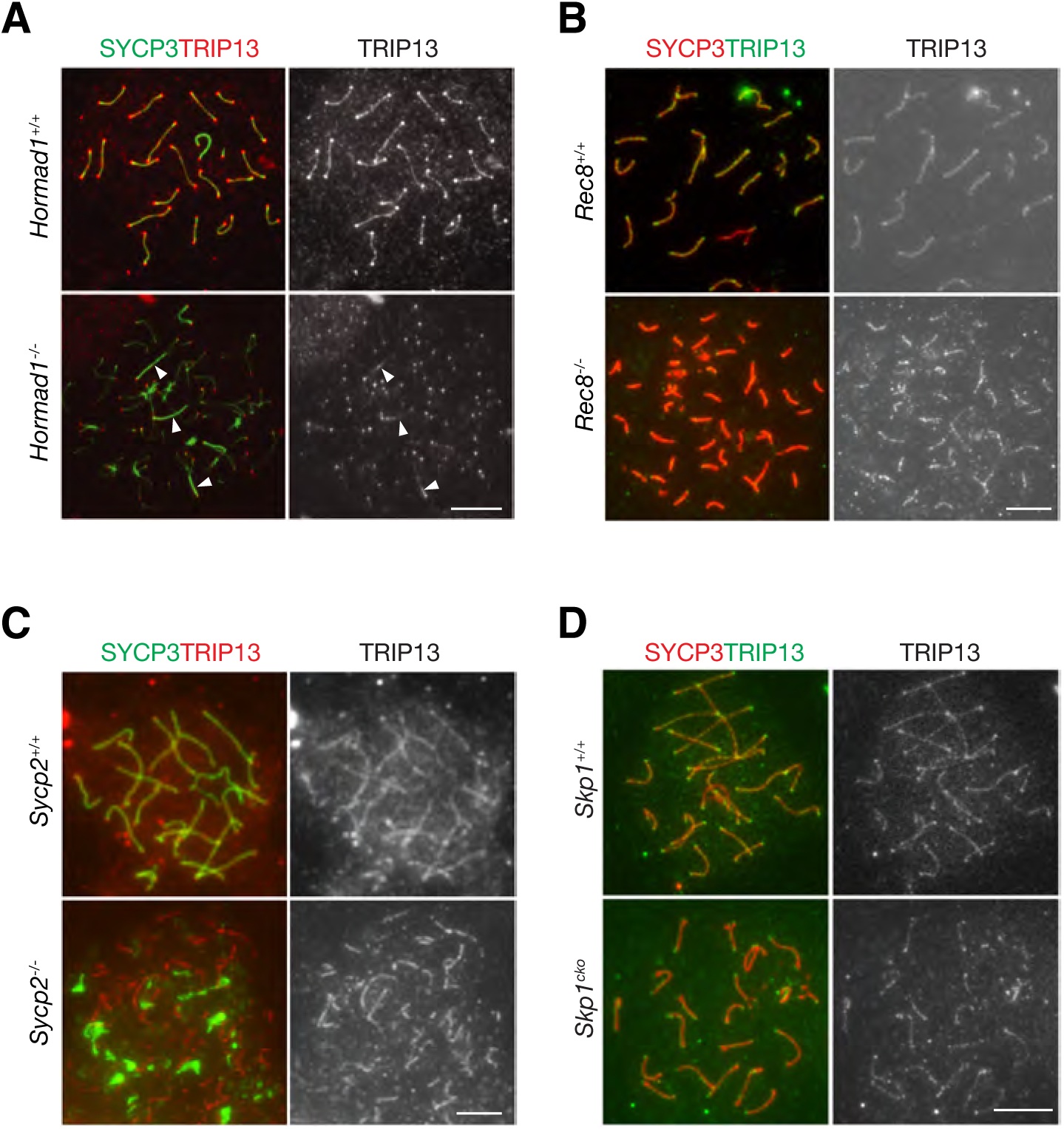
Localization of TRIP13 to the SC is independent of individual axial element components. (A) Immunofluorescent analysis of TRIP13 in *Hormad1*^-/-^ spermatocytes from 2-month-old mice. (B) Immunofluorescent analysis of TRIP13 in *Rec8*^-/-^ spermatocytes from 2-month -old mice. (C) Immunofluorescent analysis of TRIP13 in *Sycp2*^-/-^ spermatocytes from 2-month -old mice. (D) Immunofluorescent analysis of TRIP13 in *Skp1*^cKO^ spermatocytes from 2-month -old mice. Scale bars, 10 µm.

REC8, a meiosis-specific cohesin, promotes synapsis between homologs and inhibits synapsis between sister chromatids (Xu et al., 2005). In the absence of REC8, synapsis occurs between sister chromatids rather than homologous chromosomes as previously shown by super-resolution imaging (Guan et al., 2020). In *Rec8*^-/-^ spermatocytes, TRIP13 localized to synapsed sister chromatids (Figure 6B). To further probe TRIP13 recruitment to the SC, a mouse line expressing a truncated SYCP2 was used (Yang et al., 2006). SYCP2 and SYCP3 are essential components of the axial/lateral elements of the SC and interact with each other. The mutant used in this study (referred to as *Sycp2*^-/-^) expresses a truncated SYCP2 protein that lacks the C-terminal coiled-coil domain necessary for binding to SYCP3. Thus, SYCP3 failed to localize to the axial elements and formed aggregates in *Sycp2*^-/-^ spermatocytes (Figure 6C). Nevertheless, *Sycp2*^-/-^ spermatocytes formed short stretches of synapsis as previously demonstrated by electron microcopy and SYCP1 immunofluorescence (Yang et al., 2006). TRIP13 localized as filaments in *Sycp2*^-/-^ spermatocytes, strongly suggesting that its localization is independent of SYCP3 and the C-terminus of SYCP2 (Figure 6C).

Previous work has shown that SKP1, a key component of the SKP1, Cullin, F-box (SCF) complex E3 ligase, is important for HORMAD removal and chromosomal synapsis (Guan et al., 2020; Guan et al., 2022). SKP1 localizes to synapsed regions in meiotic germ cells and specifically to the lateral elements in the SC (Guan et al., 2020). We found that TRIP13 still localized to the synapsed regions in *Skp1*^cKO^ (conditional knockout) spermatocytes (Figure 6D). Taken together, these results show that the SC localization of TRIP13 is independent of HORMAD1, REC8, SYCP2, SYCP3, and SKP1, which all localize to the SC lateral elements. Such a finding is consistent with the localization of TRIP13 to the SC central region (Figure 1D).

### FLAG-tagged TRIP13 proteins are functional

In order to investigate the mechanism of TRIP13 recruitment to the SC, we generated 3×FLAG-TRIP13 (N-terminal tag) and TRIP13-3×FLAG (C-terminal tag) mice through the CRISPR/Cas9-medidated genome editing approach (Figure 7 - figure supplement 1). Two different types of tagged mice were generated in case that one fusion protein is not functional. Both alleles were transmitted through the germline of the founder mice. Western blot showed that TRIP13 existed in two isoforms in wild type (non-tagged) testis (Figure 7A). The major TRIP13 isoform was 50 kDa. The minor isoform was slightly larger than 50 kDa. Western blot analysis showed that the FLAG tagged TRIP13 fusion proteins were present in both FLAG-tagged testes. Both FLAG-tagged TRIP13 fusion proteins also existed in two isoforms in the homozygous testes, suggesting that they corresponded to the two wild type isoforms. The 3×FLAG-TRIP13 proteins were apparently slightly larger, possibly due to the linker in the N-terminally tagged proteins. The nature and physiological significance of these two isoforms were not clear, but could be due to alternative splicing or post-translational modification. Immunofluorescence analysis using anti-FLAG antibody showed that the FLAG-tagged TRIP13 proteins, like wild type TRIP13, localized to both telomeres and synaptonemal complex in pachytene spermatocytes from both FLAG/FLAG homozygous testes (Figure 7B). Importantly, both N- and C-terminally tagged homozygous mice were fertile. These results demonstrate that both N- and C-terminal tagged TRIP13 proteins localize normally and are functional.

**Figure 7.**
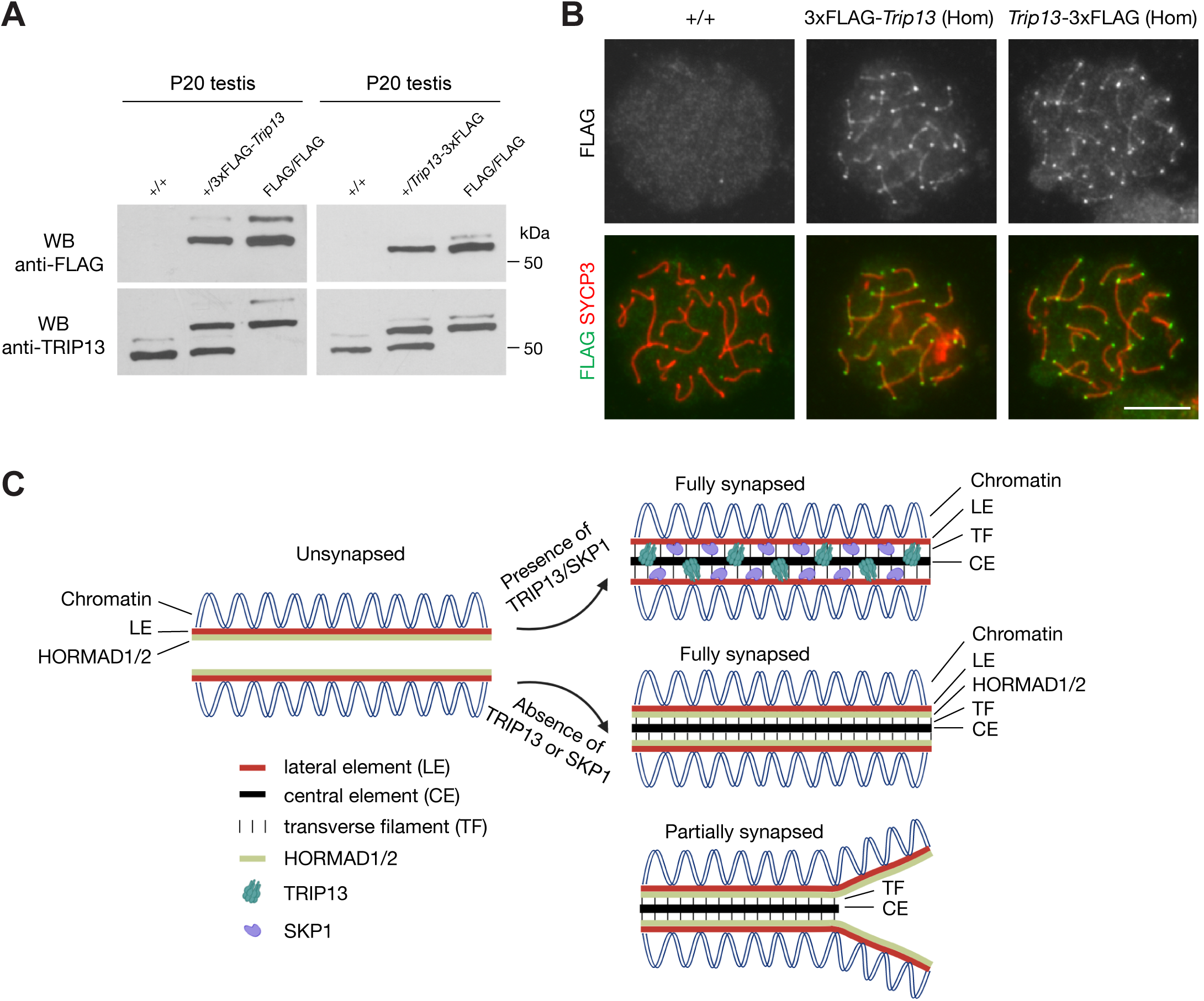
FLAG-tagged TRIP13 proteins localize correctly and are functional. (A) Western blot analysis of tagged and untagged TRIP13 proteins in testes from P20 wild type (no tag), heterozygous tagged, and homozygous tagged males. (B) Immunofluorescence of FLAG-tagged TRIP13 in pachytene spermatocytes from P20 homozygous testes. N-terminal tag, 3×FLAG-*Trip13*; C-terminal tag, *Trip13*-3×FLAG. Scale bar, 10 μm. (C) A schematic illustration of TRIP13, SKP1, HORMAD1/2, and the synaptonemal complex. Relative locations of TRIP13 and SKP1 within the SC are depicted. HORAMD1/2 are retained in synapsed regions in *Trip13*-decifient or *Skp1*-deficient spermatocytes.

To identify TRIP13-associated proteins in testis, we performed immunoprecipitation (IP) using 3×FLAG-TRIP13, TRIP13-3×FLAG, and wild type (no tag) testicular protein extracts with anti-FLAG monoclonal antibody. The immunoprecipitated proteins were eluted with FLAG peptides and subjected to mass spectrometry for protein identification. As expected, TRIP13 had more peptides in tagged TRIP13 IP than wild type (Table 1). HORMAD2, a known TRIP13 substrate protein, was also enriched in tagged TRIP13 IP (Table 1). Intriguingly, a large number of RNA-binding proteins involved in RNA splicing were highly enriched in tagged TRIP13 IP: DDX46, PUF60, RBM25, RBM39, U2AF1, U2AF2, and SRSF11 (Table 1). The biological relevance of these RNA-binding proteins to TRIP13 function remains unknown. We cannot exclude the possibility that identification of these RNA splicing factors could be immunoprecipitation artifacts.

**Table 1.** List of proteins from testis identified by co-immunoprecipitation and mass spectrometry.

| Protein | Number of peptides in anti-FLAG IP |  |  | M.W. (kD) |
| --- | --- | --- | --- | --- |
|  | 3×FLAG-TRIP13<br>testis | TRIP13-3×FLAG<br>testis | Wild type testis |  |
| TRIP13 | 28 | 36 | 4 | 51 |
| HORMAD2 | 21 | 10 | 7 | 39 |
| DDX46 | 32 | 6 | 0 | 117 |
| PUF60 | 24 | 14 | 0 | 59 |
| RBM25 | 19 | 3 | 2 | 100 |
| RBM39 | 20 | 10 | 1 | 59 |
| U2AF1 | 11 | 4 | 1 | 28 |
| U2AF2 | 15 | 9 | 3 | 54 |
| SRSF11 | 9 | 4 | 0 | 53 |
| SPATA5 | 26 | 27 | 7 | 97 |
| AP3D1 | 17 | 6 | 2 | 135 |
| PRPF40A | 16 | 3 | 0 | 108 |
| UGP2 | 15 | 3 | 0 | 57 |

## Discussion

TRIP13 and its orthologue Pch2 play an evolutionarily conserved role in depletion of meiosis-specific HORMA domain proteins from the synaptonemal complex in many species including yeast, plant, and mouse. In meiocytes, HORMA domain proteins are associated with unsynapsed chromosome axes and become depleted from the SC upon synapsis (Figure 7C). In mice with *Trip13* hypomorphic gene trap alleles, HORMAD1 and HORMAD2 persist on the synapsed SCs at the pachytene stage (Wojtasz et al., 2009). In this study, we confirmed the abnormal persistence of HORMAD1 and HORMAD2 in *Trip13*-null spermatocytes (Figure 7C). In budding yeast, deficiency of Pch2 leads to accumulation of Hop1 on the SC (Joshi et al., 2009; Subramanian et al., 2016). In *Arabidopsis*, Pch2 remodels the HORMA domain protein ASY1 (Balboni et al., 2020; Yang et al., 2020). Pch2 has been shown to localize to synapsed SC in budding yeast (Joshi et al., 2009), C. elegans (Deshong et al., 2014; Russo et al., 2023), *Arabidopsis* (Lambing et al., 2015), and rice (Miao et al., 2013), but not in mice. In mouse meiocytes, TRIP13 was known to localize to telomeres (Gomez et al., 2019). In addition to telomeres, we find that TRIP13 localizes to the SC in early pachytene spermatocytes in mice. In particular, TRIP13 localizes as a single filament between the SC lateral elements. Therefore, TRIP13/Pch2 and HORMA domain proteins have mutually exclusive localization patterns on the meiotic chromosome axes, providing an explanation for the depletion of HORMA domain proteins from the synapsed chromosomes in diverse organisms.

The transverse filaments (TF) are essential for recruitment of Pch2 to the SC. In a budding yeast Zip1 (encoding the TF component) allele with four amino acid substitutions, Pch2 is absent from the SC (Subramanian et al., 2016). In *C. elegans*, the TF component SYP-1 is essential for localization of Pch-2 on the paired chromosomes (Deshong et al., 2014). In rice, CRC1 (Pch2 orthologue) and CEP1 (TF protein) localize to the SC in an inter-dependent manner (Miao et al., 2013). In *Arabidopsis*, ZYP1 (TF component) recruits Pch2 to the SC (Yang et al., 2022). In the yeast two-hybrid assay, rice CRC1 interacts with CEP1, *Arabidopsis* ZYP1 interacts with Pch2, and mouse SYCP1 (TF protein) interacts with TRIP13 (Miao et al., 2013; Yang et al., 2022). Unlike in other species, loss of SYCP1 (TF protein) in mouse leads to a complete failure in chromosomal synapsis and thus its role in recruitment of TRIP13 to the synapsed SC can not be directly addressed (de Vries et al., 2005). Our findings on the localization of mouse TRIP13 to the SC on the sister chromatids in *Rec8*-deficient spermatocytes and the short SC stretches in *Sycp2* mutant spermatocytes demonstrate that TRIP13 recruitment is independent of SC axial element proteins REC8 and SYCP2 but support the possibility that SYCP1 could recruit TRIP13 to the synapsed SC. These findings lead to the following model: the TF protein recruits Pch2/TRIP13 to the SC upon synapsis, which in turn evicts HORMA domain proteins from the synapsed regions (Figure 7C).

Both TRIP13 and SKP1 are required for removal of HORMA domain proteins from the synapsed SC (Guan et al., 2020; Wojtasz et al., 2009). However, the relationship of TRIP13 and SKP1 is unknown. SKP1 localizes to the synapsed SC more extensively than TRIP13. While TRIP13 is only detected on the synapsed SC in early pachytene spermatocytes (Figure 1), SKP1 localizes to the synapsed SC in zygotene, all stages of pachytene, and diplotene spermatocytes (Guan et al., 2020). The localization of TRIP13 and SKP1 within the SC is different: TRIP13 is on CE/TF regions but SKP1 on LEs. While the global level of TRIP13 is reduced in *Skp1*-deficient testes, the localization of TRIP13 and SKP1 to the synapsed SC is independent (Figure 6D) (Guan et al., 2020). The SCF ubiquitin E3 ligase complex targets HORMAD1 for ubiquitination and degradation in transfected HEK293T cells (Guan et al., 2022). Loss of TRIP13 leads to accumulation of HORMAD proteins only to the synapsed SC in pachytene cells. In contrast, depletion of SKP1 causes accumulation of HORMA domain proteins, particularly HORMAD1, on both unsynapsed and synapsed chromosome axes from leptotene through diplotene meiocytes. Therefore, TRIP13 and SKP1 might regulate different pools of HORMAD proteins in meiocytes (Figure 7C).

In many organisms including mouse and human, centromeres are the last regions to synapse (Bisig et al., 2012; Brown et al., 2005; Qiao et al., 2012). The reason for this is unknown, but it might be related to the necessity to suppress deleterious non-homologous recombination at centromere and pericentromeric regions, which consist of minor and major satellite repeats respectively. The centromere was proposed to exert an inhibitory effect on synapsis in human cells (Brown et al., 2005). The inhibition needs to be relieved to achieve full synapsis, since unsynapsis at centromeres is expected to trigger the meiotic checkpoint, leading to meiotic arrest. To date, only two mouse mutants (*Skp1* and *Trip13*) exhibit unsynapsis preferentially at the centromeric end in pachytene-like cells. In addition, only these two mouse mutants display abnormal accumulation of HORMAD proteins on the synapsed SC. Intriguingly, TRIP13 localizes to telomeres in both spermatocytes and oocytes. The centromere is close to one of the telomeres in mouse. Thus, the preferential centromeric end asynapsis in *Trip13* or *Skp1*-deficient meiocytes could be related to abnormal persistence of HORMAD proteins. However, the underlying molecular mechanism warrants further investigation.

We find that TRIP13 is a dosage-dependent regulator of meiosis. The *Trip13*^+/-^ mice displayed reduced testis weight, reduced sperm count, and meiotic defects but these defects were less severe than the *Trip13*^-/-^ mice. The disease phenotypes in humans also appear to be influenced by the *TRIP13* dosage (Yost et al., 2017; Zhang et al., 2020). The dosage-dependent phenotypic variation has been reported in other mouse meiotic mutants such as *Tex11*, *Meiob*, and *Rnf212*. TEX11 localizes to meiotic chromosomes as foci and regulates crossover formation and chromosome synapsis (Yang et al., 2008). The TEX11 protein levels correlate with genome-wide recombination rates in mice (Yang et al., 2015). MEIOB, a meiosis-specific ssDNA-binding protein, is essential for meiotic recombination (Luo et al., 2013; Souquet et al., 2013). In mice with different combinations of *Meiob* alleles, MEIOB protein levels directly correlate with the severity of meiotic defects (Guo et al., 2020). Sequence variants in human RNF212 are associated with variations in genome-wide recombination rates (Chowdhury et al., 2009; Kong et al., 2008). In mouse, RNF212 forms foci on meiotic chromosomes and regulates crossover formation in a dosage-dependent manner (Reynolds et al., 2013). Therefore, dosage sensitivity appears to be a common feature of many regulators of meiosis.

## Materials and Methods

### Ethics statement

Mice were maintained and used for experimentation according to the protocol approved by the Institutional Care and Use Committee of the University of Pennsylvania.

### Mouse strains

The cryopreserved sperm from *Trip13*^+/-^ males were obtained from the Mutant Mouse Resource and Research Center (MMRCC) at UC Davis (*Trip13*^tm1.1(KOMP)Vlcg/JMmucd^, MMRRC_050223-UCD). Genotyping of wild type and knockout *Trip13* alleles was performed by separate PCR reactions using tail genomic DNA. Other mutant mouse lines used in this study were previously generated: *Hormad1*, *Sycp2*, and *Rec8* (Guan et al., 2020; Shin et al., 2010; Yang et al., 2006). Genotyping PCR primer sequences are listed in Table 2.

**Table 2.**
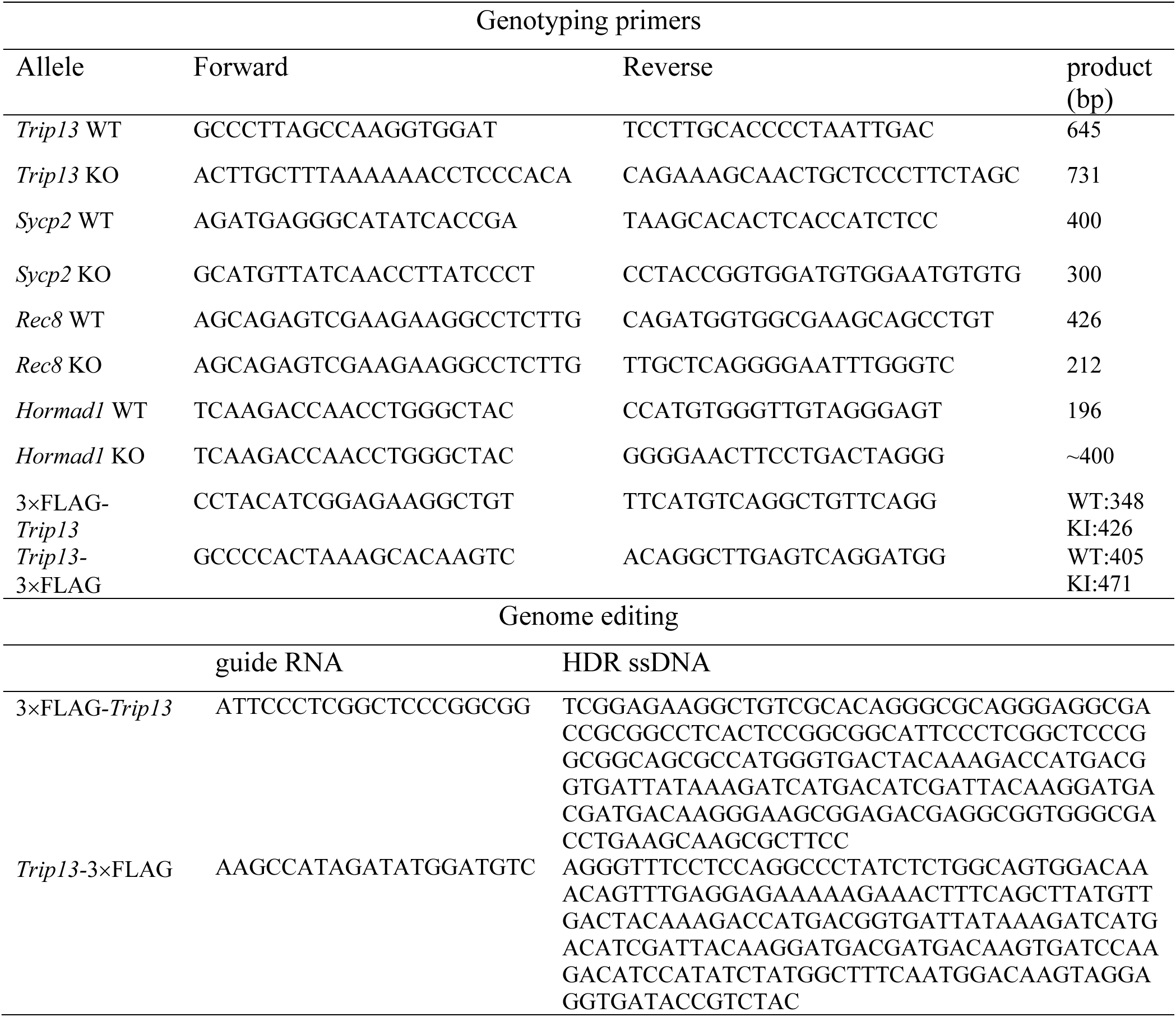
Sequences of genotyping PCR primers, sgRNA, and ssDNA templates.

### Generation of 3×FLAG-*Trip13* and *Trip13*-3×FLAG knockin mouse strains

The 3×FLAG-knockin mouse strains were generated by the CRISPR-Cas9 mediated genome editing approach. To generate N-terminally tagged knockin mice, one single-guide RNA (sgRNA) was designed to target the first exon of the mouse *Trip13* (Figure 7 – figure supplement 1A). A ssDNA repair template was created using the sequence surrounding the start codon as 5′ and 3′ homology arms. The knockin template itself included the sequence for the 3×FLAG epitope and a three-residue linker sequence. The C-terminally tagged strain was generated similarly (Figure 7 – figure supplement 1B). The guide RNA targeted exon 13 containing the stop codon. The ssDNA template contained the FLAG-encoding sequences and homology sequences flanking the insertion site. sgRNA sequences and ssDNA template sequences were shown in Table 2.

For the sgRNA, the oligo was phosphorylated, annealed, and cloned to PX330 plasmid (Addgene, Waterton, MA). After *in vitro* transcription with the MEGAshortscript T7 Kit (AM1354, Invitrogen) and purification with the MEGAclear Transcription Clean-Up Kit (AM1908, Invitrogen), a mixture of Cas9 mRNA [100ng/µl; Trilink, catalog number L-7206) + 0.5 µl of the sgRNA (35 ng/µl) + 100ng/ul of ssDNA template] was prepared and injected into zygotes. The injected zygotes were cultured in KSOM medium at 37°C in a 5% CO_2_ incubator until the two-cell stage. The two-cell embryos were transferred into oviducts of 0.5-day post-coitum pseudopregnant ICR foster mothers. Founder mice were bred to wild type mice to obtain germline transmission. The N-terminal 3×FLAG allele and the C-terminal 3×FLAG allele were PCR amplified and sequenced to confirm the insertion. PCR genotyping primers are listed in Table 2.

### Production of anti-MAJIN antibodies

The short isoform of mouse MAJIN (amino acids 1-124; XM_036161650.1 and XP_036017543.1) was expressed as a 6xHis-MAJIN recombinant protein in *E. coli* using the pQE-30 vector. The recombinant protein was affinity purified with the Ni-NTA agarose. Two guinea pigs were immunized at Cocalico Biologicals Inc. (Reamstown, PA), resulting in two antisera (UP-GP140 and UP-GP141). Both anti-sera were used for immunofluorescence of nuclear spreads of spermatocytes.

### Histological, immunofluorescence, and surface nuclear spread analyses

For histology, testes or ovaries were fixed in Bouin’s solution at room temperature overnight, embedded with paraffin, and sectioned at 8 μm. Sections were stained with hematoxylin and eosin. For immunofluorescence analysis, testes were fixed in 4% paraformaldehyde (in 1×PBS) overnight at 4°C, dehydrated in 30% sucrose (in 1×PBS) overnight, and sectioned at 8 μm in a cryostat. Surface nuclear spread analysis was described before (Peters et al., 1997). Briefly, testicular tubules or ovarian tissues were soaked in hypotonic treatment buffer (30 mM Tris, 50 mM sucrose, 17 mM trisodium citrate dihydrate, 5 mM EDTA, 0.5 mM DTT, 1 mM PMSF). Then the cells were suspended in 100 mM sucrose and spread by physical disruption on “PTFE” printed slides that were previously soaked with paraformaldehyde solution containing Triton X-100 and sodium borate. Antibodies used for immunostaining are listed in Table 3.

**Table 3.** List of primary antibodies.

| Reagent | Dilution | Source | Catalog # or reference |
| --- | --- | --- | --- |
| Mouse anti-ACTB mab | 1:2000 | Sigma | Cat# A5441; RRID: AB_476744 |
| Human anti-centromere (CREST) | 1:500 | Antibodies Incorporated | Cat# 15-234; RRID: AB_2687472 |
| Rabbit anti-SYCP1 | 1:300 | Abcam | Cat# ab15090; RRID: AB_301636 |
| Guinea pig anti-SYCP2 | 1:150 | Custom made | (Yang et al., 2006) |
| Mouse anti-SYCP3 mab | 1:500 | Abcam | Cat# ab97672; RRID: AB_10678841 |
| Rabbit anti-SYCP3 | 1:500 IF; 1:2000 WB | ProteinTech Group | Cat# 23024-1-AP; RRID: AB_11232426 |
| Rabbit anti-HORMAD1 | 1:300 | ProteinTech Group | Cat# 13917-1-AP; RRID: AB_2120844 |
| Rabbit anti-HORMAD2 | 1:500 | Custom made, a gift from Toth lab | (Wojtasz et al., 2009) |
| Rabbit anti-REC114 | 1:100 | Custom made, a gift from Keeney Lab | (Boekhout et al., 2019) |
| Rabbit anti-REC8 | 1:200 | Custom made, a gift from Mengcheng Luo lab | N/A |
| Mouse anti-FLAG mab | 1:300IF; 1:4000WB | Sigma | Cat# F3165, RRID: AB_259529 |
| Rabbit anti-SKP1 | 1:200 IF; 1:1000 WB | Cell Signaling | Cat# 12248S, RRID: AB_2754993 |
| Rabbit anti-TRIP13 | 1:150 IF; 1:1000 WB | ProteinTech Group | 19602-1-AP |
| Rabbit anti-SYCE1 | 1:200 IF; 1:1000 WB | Custom made | N/A |
| Mouse anti-SYCP1 mab | 1:200 IF; 1:1000 WB | Custom made, 1:200; a gift from Chritster Hoog lab | N/A |
| Rabbit anti-CENP-C | 1:500 | Custom made, a gift from Y. Watanabe | (Kim et al., 2015) |
| Mouse anti-TRF1 mab | 1:100 | Sigma | T1948 |
| Guinea pig anti- MAJIN | 1:100 | Custom made; this study | N/A |

### Imaging

Histological images were captured on the Leica DM5500B microscope with a DFC450 digital camera (Leica Microsystems, Wetzlar, Germany). Most immunolabeled chromosome spread images were taken on the Leica DM5500B microscope with an ORCA Flash4.0 digital monochrome camera (Hamamatsu Photonics, Bridgewater, NJ). Confocal microscopy of immunolabeled chromosome spreads was performed on a Leica SP5 II confocal (Leica Microsystems, Wetzlar, Germany) with an 100× (1.46 NA) oil immersion objective lens. Images were deconvolved with Huygens Essential deconvolution software (Scientific Volume Imaging B.V., Hilversum, Netherlands).

### Western blot analysis

Testes were homogenized in the lysis buffer (50 mM Tris-HCl, pH 8.0, 150 mM NaCl, 1% Trion X-100, 0.5% sodium deoxycholate, 5 mM MgCl_2_, and 1 mM DTT supplemented with 1 mM PMSF). 40 μg of protein samples were resolved by SDS-PAGE, transferred onto PDVF membranes, and immunoblotted with primary antibodies (Table 3).

## Acknowledgements

We thank Gordon Ruthel at the PennVet Imaging Core for help with super-resolution microscopy, Hsin Yao Tang and Thomas Beer at Wistar Proteomics Core for help with mass spectrometry, Christer Hoog for SYCP1 antibody, Attila Toth for HORMAD2 antibody, Scott Keeney for anti-REC114 antibody, Yoshinori Watanabe and Takashi Akera for CENP-C antibody.

---

**Figure 1 - figure supplement 1.**
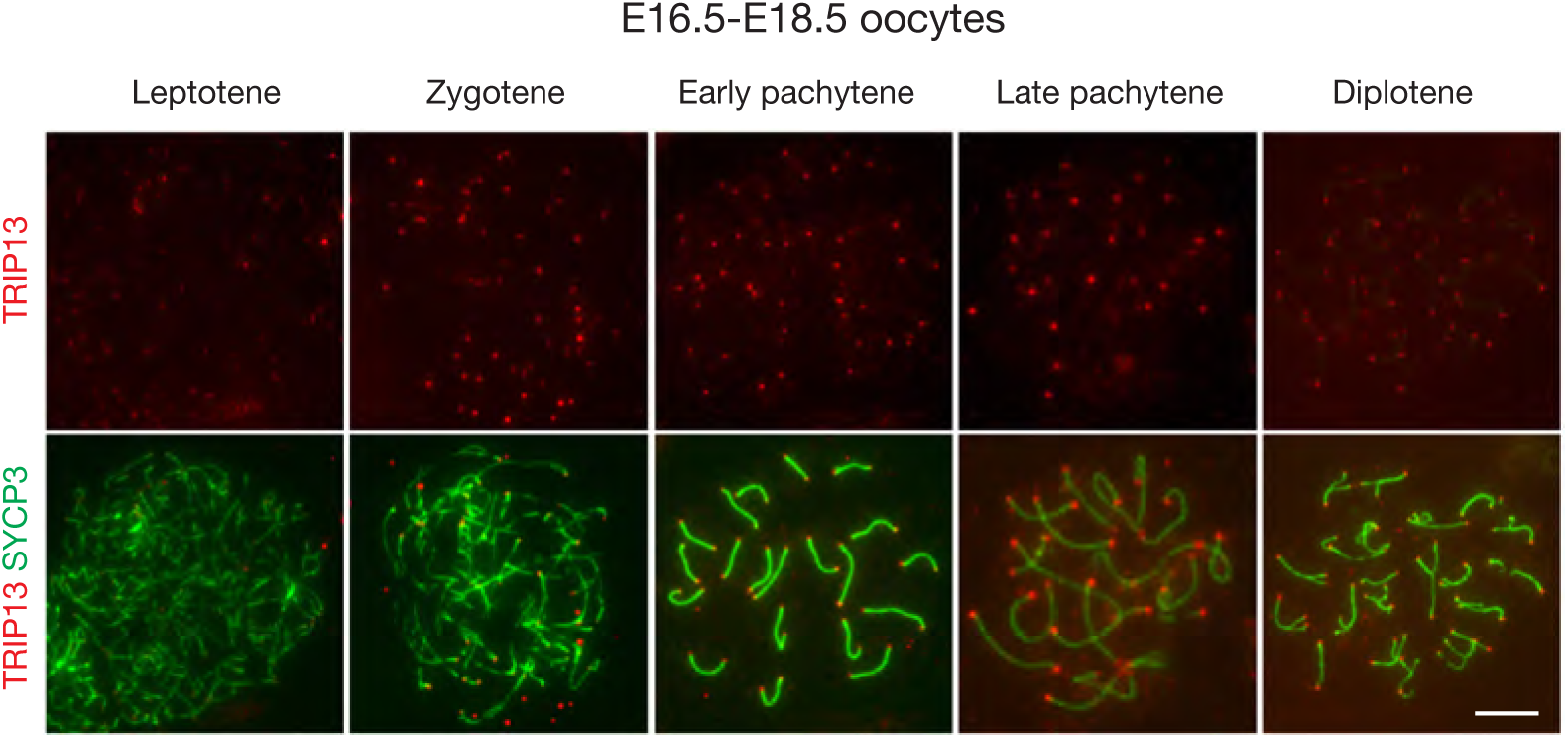
Immunofluorescent analysis of TRIP13 in spread nuclei of oocytes from embryonic day 16.5 (E16.5) and E18.5 female embryos. Scale bars, 10 µm.

**Figure 2 - figure supplement 1.**
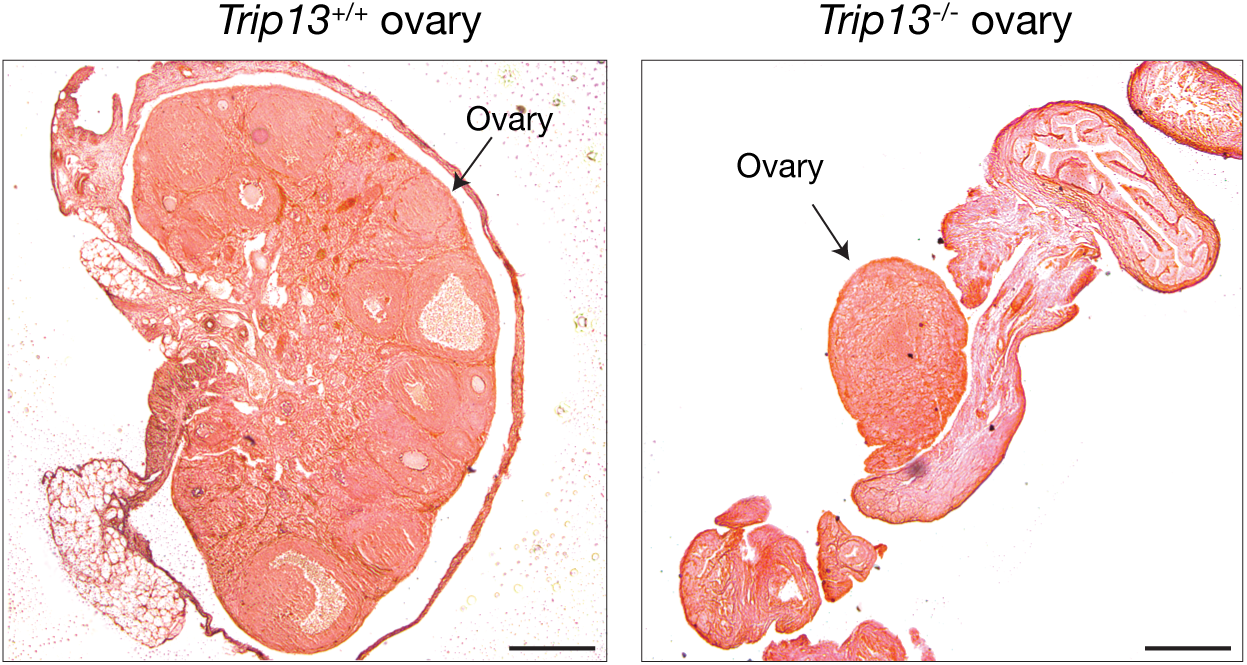
Histological analysis of ovaries from adult (8-week) wild type and *Trip13*^-/-^ females. Scale bars, 200 µm.

**Figure 4 - figure supplement 1.**
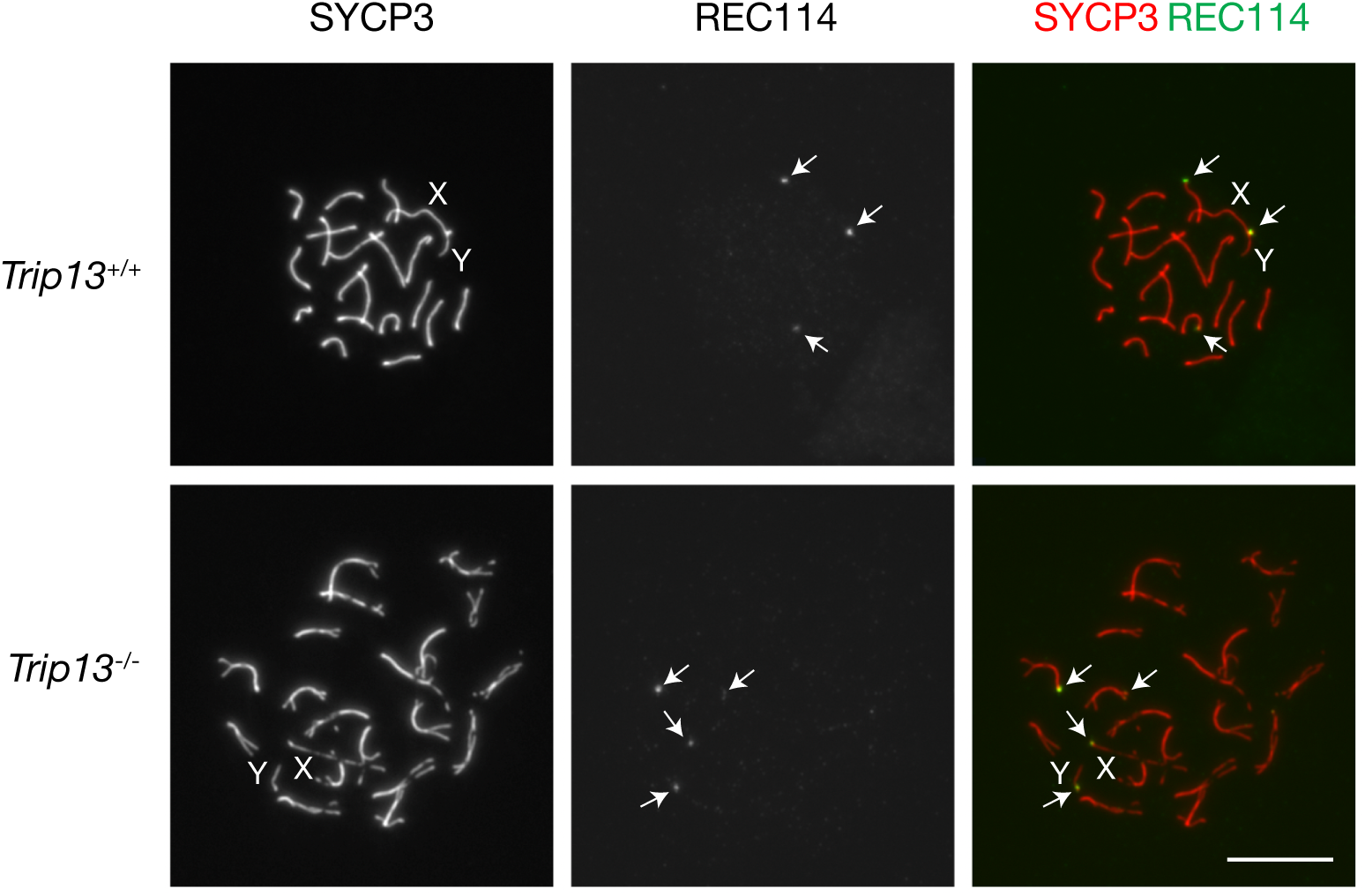
Immunofluorescent analysis of REC114 in spread nuclei of spermatocytes from P28 wild type (*Trip13*^+/+^) and *Trip13*^-/-^ testes. Wild type, early pachytene; *Trip13*^-/-^, pachytene-like. REC114 foci are indicated by arrows. Scale bar, 10 μm.

**Figure 7 - figure supplement 1.**
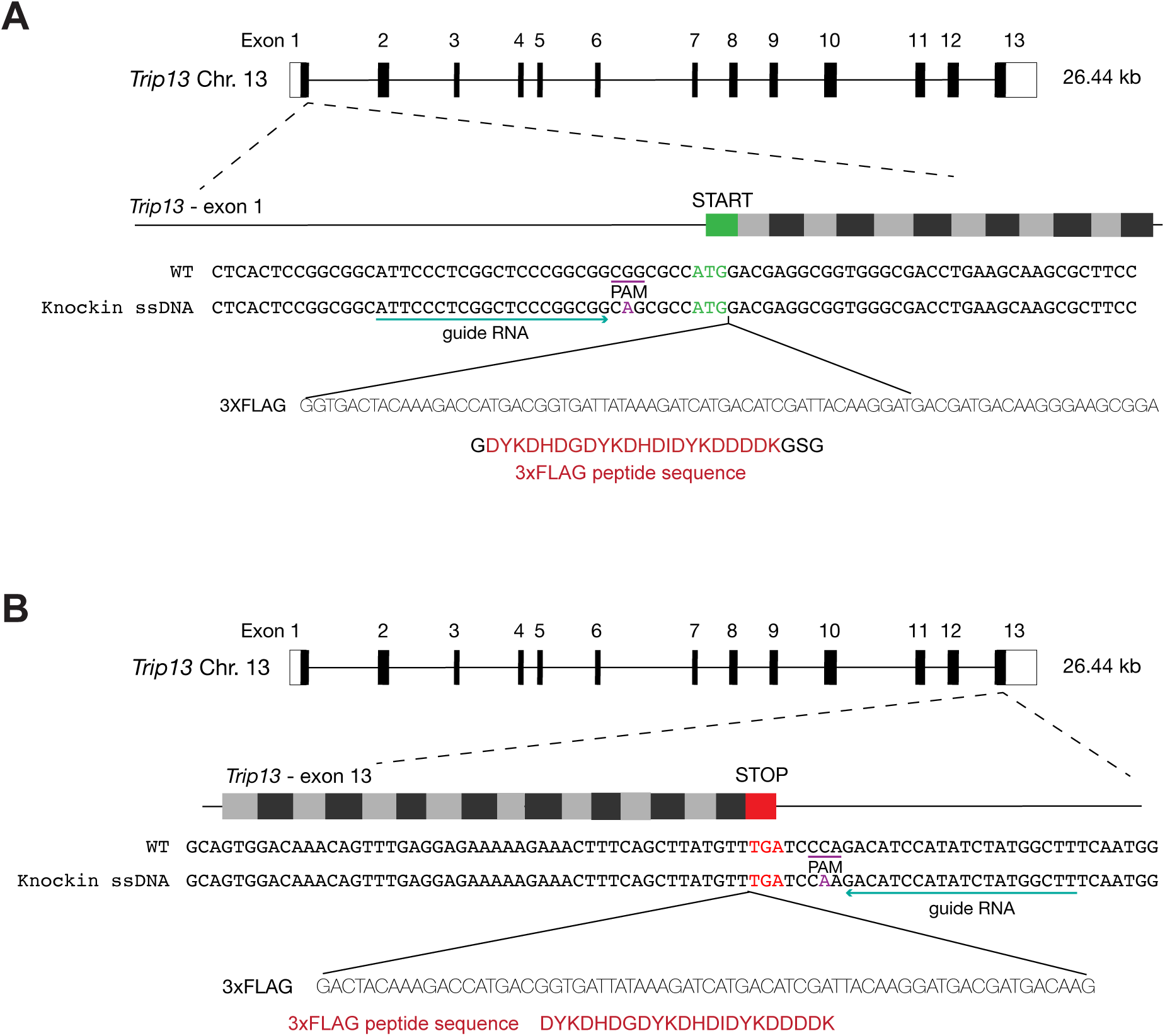
Generation of two *Trip13* FLAG tagged mouse lines. (A) Illustration of the *Trip13* gene structure with the 3×FLAG tag at the N terminus. The guide RNA sequence is underlined in green. The single strand DNA (ssDNA) oligo template (200 nt) contains the 3×FLAG-encoding sequence. The PAM site is underlined in purple. One base in the PAM site is mutated in the ssDNA oligo template to prevent cutting of the template strand (in purple). The position of 3×FLAG in-frame insertion is designated (^) and occurs just after the endogenous start codon. Filled bars, coding regions; Open bars, 5′ or 3′ UTRs. (B) Illustration of the *Trip13* gene structure with the 3×FLAG tag at the C terminus. The PAM cut site and guide RNA were on the reverse strand, but for clarity, the forward strand sequence is shown. The 3×FLAG was inserted just before the endogenous stop codon.

## References

1. Balboni, M., Yang, C., Komaki, S., Brun, J. and Schnittger, A. (2020). COMET Functions as a PCH2 Cofactor in Regulating the HORMA Domain Protein ASY1. Curr Biol 30, 4113–4127 e4116.

2. Bisig, C. G., Guiraldelli, M. F., Kouznetsova, A., Scherthan, H., Hoog, C., Dawson, D. S. and Pezza, R. J. (2012). Synaptonemal complex components persist at centromeres and are required for homologous centromere pairing in mouse spermatocytes. PLoS genetics 8, e1002701.

3. Boekhout, M., Karasu, M. E., Wang, J., Acquaviva, L., Pratto, F., Brick, K., Eng, D. Y., Xu, J., Camerini-Otero, R. D., Patel, D. J., et al. (2019). REC114 Partner ANKRD31 Controls Number, Timing, and Location of Meiotic DNA Breaks. Molecular cell 74, 1053–1068.e1058.

4. Brown, P. W., Judis, L., Chan, E. R., Schwartz, S., Seftel, A., Thomas, A. and Hassold, T. J. (2005). Meiotic synapsis proceeds from a limited number of subtelomeric sites in the human male. American Journal of Human Genetics 77, 556–566.

5. Chen, C., Jomaa, A., Ortega, J. and Alani, E. E. (2014). Pch2 is a hexameric ring ATPase that remodels the chromosome axis protein Hop1. Proceedings of the National Academy of Sciences of the United States of America 111, 44.

6. Chowdhury, R., Bois, P. R., Feingold, E., Sherman, S. L. and Cheung, V. G. (2009). Genetic analysis of variation in human meiotic recombination. PLoS genetics 5, e1000648.

7. Daniel, K., Lange, J., Hached, K., Fu, J., Anastassiadis, K., Roig, I., Cooke, H. J., Stewart, A. F., Wassmann, K., Jasin, M., et al. (2011). Meiotic homologue alignment and its quality surveillance are controlled by mouse HORMAD1. Nature cell biology 13, 599–610.

8. de Vries, F. A., de Boer, E., van den Bosch, M., Baarends, W. M., Ooms, M., Yuan, L., Liu, J. G., van Zeeland, A. A., Heyting, C. and Pastink, A. (2005). Mouse Sycp1 functions in synaptonemal complex assembly, meiotic recombination, and XY body formation. Genes & development 19, 1376–1389.

9. Deshong, A. J., Ye, A. L., Lamelza, P. and Bhalla, N. (2014). A quality control mechanism coordinates meiotic prophase events to promote crossover assurance. PLoS Genet 10, e1004291.

10. Fukuda, T., Daniel, K., Wojtasz, L., Toth, A. and Hoog, C. (2010). A novel mammalian HORMA domain-containing protein, HORMAD1, preferentially associates with unsynapsed meiotic chromosomes. Experimental cell research 316, 158–171.

11. Gomez, H. L., Felipe-Medina, N., Condezo, Y. B., Garcia-Valiente, R., Ramos, I., Suja, J. A., Barbero, J. L., Roig, I., Sanchez-Martin, M., de Rooij, D. G., et al. (2019). The PSMA8 subunit of the spermatoproteasome is essential for proper meiotic exit and mouse fertility. PLoS Genet 15, e1008316.

12. Guan, Y., Leu, N. A., Ma, J., Chmatal, L., Ruthel, G., Bloom, J. C., Lampson, M. A., Schimenti, J. C., Luo, M. and Wang, P. J. (2020). SKP1 drives the prophase I to metaphase I transition during male meiosis. Science advances 6, eaaz2129.

13. Guan, Y., Lin, H., Leu, N. A., Ruthel, G., Fuchs, S. Y., Busino, L., Luo, M. and Wang, P. J. (2022). SCF ubiquitin E3 ligase regulates DNA double-strand breaks in early meiotic recombination. Nucleic Acids Res 50, 5129–5144.

14. Guo, R., Xu, Y., Leu, N. A., Zhang, L., Fuchs, S. Y., Ye, L. and Wang, P. J. (2020). The ssDNA-binding protein MEIOB acts as a dosage-sensitive regulator of meiotic recombination. Nucleic acids research 48, 12219–12233.

15. Handel, M. A. and Schimenti, J. C. (2010). Genetics of mammalian meiosis: regulation, dynamics and impact on fertility. Nature reviews.Genetics 11, 124–136.

16. Herruzo, E., Lago-Maciel, A., Baztan, S., Santos, B., Carballo, J. A. and San-Segundo, P. A. (2021). Pch2 orchestrates the meiotic recombination checkpoint from the cytoplasm. PLoS Genet 17, e1009560.

17. Herruzo, E., Ontoso, D., Gonzalez-Arranz, S., Cavero, S., Lechuga, A. and San-Segundo, P. A. (2016). The Pch2 AAA+ ATPase promotes phosphorylation of the Hop1 meiotic checkpoint adaptor in response to synaptonemal complex defects. Nucleic Acids Res 44, 7722–7741.

18. Hunter, N. (2015). Meiotic Recombination: The Essence of Heredity. Cold Spring Harbor perspectives in biology 7, 10.1101/cshperspect.a016618.

19. Joshi, N., Barot, A., Jamison, C. and Borner, G. V. (2009). Pch2 links chromosome axis remodeling at future crossover sites and crossover distribution during yeast meiosis. PLoS Genet 5, e1000557.

20. Karlseder, J., Kachatrian, L., Takai, H., Mercer, K., Hingorani, S., Jacks, T. and de Lange, T. (2003). Targeted deletion reveals an essential function for the telomere length regulator Trf1. Molecular and cellular biology 23, 6533–6541.

21. Kim, J., Ishiguro, K., Nambu, A., Akiyoshi, B., Yokobayashi, S., Kagami, A., Ishiguro, T., Pendas, A. M., Takeda, N., Sakakibara, Y., et al. (2015). Meikin is a conserved regulator of meiosis-I-specific kinetochore function. Nature 517, 466–471.

22. Kim, Y., Rosenberg, S. C., Kugel, C. L., Kostow, N., Rog, O., Davydov, V., Su, T. Y., Dernburg, A. F. and Corbett, K. D. (2014). The chromosome axis controls meiotic events through a hierarchical assembly of HORMA domain proteins. Dev Cell 31, 487–502.

23. Klare, K., Weir, J. R., Basilico, F., Zimniak, T., Massimiliano, L., Ludwigs, N., Herzog, F. and Musacchio, A. (2015). CENP-C is a blueprint for constitutive centromere-associated network assembly within human kinetochores. The Journal of cell biology 210, 11–22.

24. Kogo, H., Tsutsumi, M., Inagaki, H., Ohye, T., Kiyonari, H. and Kurahashi, H. (2012a). HORMAD2 is essential for synapsis surveillance during meiotic prophase via the recruitment of ATR activity. Genes Cells 17, 897–912.

25. Kogo, H., Tsutsumi, M., Ohye, T., Inagaki, H., Abe, T. and Kurahashi, H. (2012b). HORMAD1-dependent checkpoint/surveillance mechanism eliminates asynaptic oocytes. Genes Cells 17, 439–454.

26. Kong, A., Thorleifsson, G., Stefansson, H., Masson, G., Helgason, A., Gudbjartsson, D. F., Jonsdottir, G. M., Gudjonsson, S. A., Sverrisson, S., Thorlacius, T., et al. (2008). Sequence variants in the RNF212 gene associate with genome-wide recombination rate. Science (New York, N.Y.) 319, 1398–1401.

27. Kwon, M. S., Hori, T., Okada, M. and Fukagawa, T. (2007). CENP-C is involved in chromosome segregation, mitotic checkpoint function, and kinetochore assembly. Mol Biol Cell 18, 2155–2168.

28. Lambing, C., Osman, K., Nuntasoontorn, K., West, A., Higgins, J. D., Copenhaver, G. P., Yang, J., Armstrong, S. J., Mechtler, K., Roitinger, E., et al. (2015). Arabidopsis PCH2 Mediates Meiotic Chromosome Remodeling and Maturation of Crossovers. PLoS Genet 11, e1005372.

29. Li, X. C. and Schimenti, J. C. (2007). Mouse pachytene checkpoint 2 (trip13) is required for completing meiotic recombination but not synapsis. PLoS genetics 3, e130.

30. Long, J., Huang, C., Chen, Y., Zhang, Y., Shi, S., Wu, L., Liu, Y., Liu, C., Wu, J. and Lei, M. (2017). Telomeric TERB1-TRF1 interaction is crucial for male meiosis. Nat Struct Mol Biol 24, 1073–1080.

31. Luo, M., Yang, F., Leu, N. A., Landaiche, J., Handel, M. A., Benavente, R., La Salle, S. and Wang, P. J. (2013). MEIOB exhibits single-stranded DNA-binding and exonuclease activities and is essential for meiotic recombination. Nature communications 4, 2788.

32. Luo, X., Tang, Z., Xia, G., Wassmann, K., Matsumoto, T., Rizo, J. and Yu, H. (2004). The Mad2 spindle checkpoint protein has two distinct natively folded states. Nat Struct Mol Biol 11, 338–345.

33. Miao, C., Tang, D., Zhang, H., Wang, M., Li, Y., Tang, S., Yu, H., Gu, M. and Cheng, Z. (2013). Central region component1, a novel synaptonemal complex component, is essential for meiotic recombination initiation in rice. Plant Cell 25, 2998–3009.

34. Pacheco, S., Marcet-Ortega, M., Lange, J., Jasin, M., Keeney, S. and Roig, I. (2015). The ATM signaling cascade promotes recombination-dependent pachytene arrest in mouse spermatocytes. PLoS genetics 11, e1005017.

35. Page, S. L. and Hawley, R. S. (2004). The genetics and molecular biology of the synaptonemal complex. Annual Review of Cell and Developmental Biology 20, 525–558.

36. Papanikos, F., Clement, J. A. J., Testa, E., Ravindranathan, R., Grey, C., Dereli, I., Bondarieva, A., Valerio-Cabrera, S., Stanzione, M., Schleiffer, A., et al. (2019). Mouse ANKRD31 Regulates Spatiotemporal Patterning of Meiotic Recombination Initiation and Ensures Recombination between X and Y Sex Chromosomes. Mol Cell 74, 1069–1085 e1011.

37. Peters, A. H., Plug, A. W., van Vugt, M. J. and de Boer, P. (1997). A drying-down technique for the spreading of mammalian meiocytes from the male and female germline. Chromosome research : an international journal on the molecular, supramolecular and evolutionary aspects of chromosome biology 5, 66–68.

38. Prince, J. P. and Martinez-Perez, E. (2022). Functions and Regulation of Meiotic HORMA-Domain Proteins. Genes (Basel) 13.

39. Qiao, H., Chen, J. K., Reynolds, A., Hoog, C., Paddy, M. and Hunter, N. (2012). Interplay between synaptonemal complex, homologous recombination, and centromeres during mammalian meiosis. PLoS genetics 8, e1002790.

40. Raina, V. B. and Vader, G. (2020). Homeostatic Control of Meiotic Prophase Checkpoint Function by Pch2 and Hop1. Curr Biol 30, 4413–4424 e4415.

41. Reynolds, A., Qiao, H., Yang, Y., Chen, J. K., Jackson, N., Biswas, K., Holloway, J. K., Baudat, F., de Massy, B., Wang, J., et al. (2013). RNF212 is a dosage-sensitive regulator of crossing-over during mammalian meiosis. Nature genetics 45, 269–278.

42. Rinaldi, V. D., Bolcun-Filas, E., Kogo, H., Kurahashi, H. and Schimenti, J. C. (2017). The DNA Damage Checkpoint Eliminates Mouse Oocytes with Chromosome Synapsis Failure. Mol Cell 67, 1026–1036 e1022.

43. Roeder, G. S. and Bailis, J. M. (2000). The pachytene checkpoint. Trends in genetics : TIG; Trends in genetics : TIG 16, 395–403.

44. Roig, I., Dowdle, J. A., Toth, A., de Rooij, D. G., Jasin, M. and Keeney, S. (2010). Mouse TRIP13/PCH2 is required for recombination and normal higher-order chromosome structure during meiosis. PLoS genetics 6, 10.1371/journal.pgen.1001062.

45. Russo, A. E., Giacopazzi, S., Deshong, A., Menon, M., Ortiz, V., Ego, K. M., Corbett, K. D. and Bhalla, N. (2023). The conserved AAA ATPase PCH-2 distributes its regulation of meiotic prophase events through multiple meiotic HORMADs in C. elegans. PLoS Genet 19, e1010708.

46. San-Segundo, P. A. and Roeder, G. S. (1999). Pch2 links chromatin silencing to meiotic checkpoint control. Cell 97, 313–324.

47. Shibuya, H., Hernandez-Hernandez, A., Morimoto, A., Negishi, L., Hoog, C. and Watanabe, Y. (2015). MAJIN Links Telomeric DNA to the Nuclear Membrane by Exchanging Telomere Cap. Cell 163, 1252–1266.

48. Shibuya, H., Ishiguro, K. and Watanabe, Y. (2014). The TRF1-binding protein TERB1 promotes chromosome movement and telomere rigidity in meiosis. Nature cell biology 16, 145–156.

49. Shin, Y. H., Choi, Y., Erdin, S. U., Yatsenko, S. A., Kloc, M., Yang, F., Wang, P. J., Meistrich, M. L. and Rajkovic, A. (2010). Hormad1 mutation disrupts synaptonemal complex formation, recombination, and chromosome segregation in mammalian meiosis. PLoS genetics 6, e1001190.

50. Sironi, L., Mapelli, M., Knapp, S., De Antoni, A., Jeang, K. T. and Musacchio, A. (2002). Crystal structure of the tetrameric Mad1-Mad2 core complex: implications of a ’safety belt’ binding mechanism for the spindle checkpoint. EMBO J 21, 2496–2506.

51. Souquet, B., Abby, E., Herve, R., Finsterbusch, F., Tourpin, S., Le Bouffant, R., Duquenne, C., Messiaen, S., Martini, E., Bernardino-Sgherri, J., et al. (2013). MEIOB Targets Single-Strand DNA and Is Necessary for Meiotic Recombination. PLoS genetics 9, e1003784.

52. Stanzione, M., Baumann, M., Papanikos, F., Dereli, I., Lange, J., Ramlal, A., Trankner, D., Shibuya, H., de Massy, B., Watanabe, Y., et al. (2016). Meiotic DNA break formation requires the unsynapsed chromosome axis-binding protein IHO1 (CCDC36) in mice. Nature cell biology 18, 1208–1220.

53. Subramanian, V. V., MacQueen, A. J., Vader, G., Shinohara, M., Sanchez, A., Borde, V., Shinohara, A. and Hochwagen, A. (2016). Chromosome Synapsis Alleviates Mek1-Dependent Suppression of Meiotic DNA Repair. PLoS Biol 14, e1002369.

54. West, A. M., Rosenberg, S. C., Ur, S. N., Lehmer, M. K., Ye, Q., Hagemann, G., Caballero, I., Uson, I., MacQueen, A. J., Herzog, F., et al. (2019). A conserved filamentous assembly underlies the structure of the meiotic chromosome axis. Elife 8.

55. West, A. M. V., Komives, E. A. and Corbett, K. D. (2018). Conformational dynamics of the Hop1 HORMA domain reveal a common mechanism with the spindle checkpoint protein Mad2. Nucleic Acids Res 46, 279–292.

56. Wojtasz, L., Cloutier, J. M., Baumann, M., Daniel, K., Varga, J., Fu, J., Anastassiadis, K., Stewart, A. F., Reményi, A., Turner, J. M., et al. (2012). Meiotic DNA double-strand breaks and chromosome asynapsis in mice are monitored by distinct HORMAD2-independent and -dependent mechanisms. Genes Dev 26, 958–973.

57. Wojtasz, L., Daniel, K., Roig, I., Bolcun-Filas, E., Xu, H., Boonsanay, V., Eckmann, C. R., Cooke, H. J., Jasin, M., Keeney, S., et al. (2009). Mouse HORMAD1 and HORMAD2, two conserved meiotic chromosomal proteins, are depleted from synapsed chromosome axes with the help of TRIP13 AAA-ATPase. PLoS genetics 5, e1000702.

58. Xu, H., Beasley, M. D., Warren, W. D., van der Horst, G. T. and McKay, M. J. (2005). Absence of mouse REC8 cohesin promotes synapsis of sister chromatids in meiosis. Developmental cell 8, 949–961.

59. Xu, H., Tong, Z., Ye, Q., Sun, T., Hong, Z., Zhang, L., Bortnick, A., Cho, S., Beuzer, P., Axelrod, J., et al. (2019). Molecular organization of mammalian meiotic chromosome axis revealed by expansion STORM microscopy. Proc Natl Acad Sci U S A 116, 18423–18428.

60. Yang, C., Hu, B., Portheine, S. M., Chuenban, P. and Schnittger, A. (2020). State changes of the HORMA protein ASY1 are mediated by an interplay between its closure motif and PCH2. Nucleic Acids Res 48, 11521–11535.

61. Yang, C., Sofroni, K., Hamamura, Y., Hu, B., Elbasi, H. T., Balboni, M., Chu, L., Stang, D., Heese, M. and Schnittger, A. (2022). ZYP1-mediated recruitment of PCH2 to the synaptonemal complex remodels the chromosome axis leading to crossover restriction. Nucleic Acids Res 50, 12924–12937.

62. Yang, F., De La Fuente, R., Leu, N. A., Baumann, C., McLaughlin, K. J. and Wang, P. J. (2006). Mouse SYCP2 is required for synaptonemal complex assembly and chromosomal synapsis during male meiosis. The Journal of cell biology 173, 497–507.

63. Yang, F., Gell, K., van der Heijden, G. W., Eckardt, S., Leu, N. A., Page, D. C., Benavente, R., Her, C., Hoog, C., McLaughlin, K. J., et al. (2008). Meiotic failure in male mice lacking an X-linked factor. Genes & development 22, 682–691.

64. Yang, F., Silber, S., Leu, N. A., Oates, R. D., Marszalek, J. D., Skaletsky, H., Brown, L. G., Rozen, S., Page, D. C. and Wang, P. J. (2015). TEX11 is mutated in infertile men with azoospermia and regulates genome-wide recombination rates in mouse. EMBO molecular medicine 7, 1198–1210.

65. Ye, Q., Kim, D. H., Dereli, I., Rosenberg, S. C., Hagemann, G., Herzog, F., Toth, A., Cleveland, D. W. and Corbett, K. D. (2017). The AAA+ ATPase TRIP13 remodels HORMA domains through N-terminal engagement and unfolding. EMBO J 36, 2419–2434.

66. Yost, S., de Wolf, B., Hanks, S., Zachariou, A., Marcozzi, C., Clarke, M., de Voer, R., Etemad, B., Uijttewaal, E., Ramsay, E., et al. (2017). Biallelic TRIP13 mutations predispose to Wilms tumor and chromosome missegregation. Nat Genet 49, 1148–1151.

67. Zhang, Z., Li, B., Fu, J., Li, R., Diao, F., Li, C., Chen, B., Du, J., Zhou, Z., Mu, J., et al. (2020). Bi-allelic Missense Pathogenic Variants in TRIP13 Cause Female Infertility Characterized by Oocyte Maturation Arrest. American Journal of Human Genetics 107, 15–23.

68. Zickler, D. and Kleckner, N. (2015). Recombination, Pairing, and Synapsis of Homologs during Meiosis. Cold Spring Harbor perspectives in biology 7, 10.1101/cshperspect.a016626.

